# Dynamic subcellular proteomics identifies novel regulators of adipocyte insulin action

**DOI:** 10.1101/2025.03.13.642412

**Authors:** Olivia J. Conway, Josie A. Christopher, Lisa M. Breckels, Hanqi Li, Dilip Menon, Lu Liang, Satish Patel, David C. Gershlick, David B. Savage, Michael P. Weekes, Kathryn S. Lilley, Daniel J. Fazakerley

## Abstract

Insulin acts on adipocytes to suppress lipolysis and increase glucose uptake to control whole-body glucose and lipid metabolism. Regulation of these processes by insulin signalling depends on changes in protein localisation. However, the extent of insulin-stimulated changes to the adipocyte spatial proteome, and the importance of these in the cellular insulin response, is unknown. Here, we use subcellular proteomics approaches to map acute insulin-stimulated protein relocalisation in adipocytes on a cell-wide scale. These data revealed extensive insulin-regulated protein redistribution, with hundreds of novel insulin-responsive proteins. These included the uncharacterised protein C3ORF18, which redistributed to the PM in response to insulin. Studies in C3ORF18-depleted adipocytes suggest this protein is required for maximal insulin signalling. Overall, our data highlight the scale of protein relocalisation in the adipocyte insulin response, and provide an accessible resource to inform further studies into how changes in protein localisation contribute to cellular insulin responses.

## Introduction

White adipose tissue is sensitive to hormonal and metabolic inputs that balance energy storage and energy release, coordinating adipocyte metabolism and organismal needs. Postprandially, increased circulating insulin promotes adipocyte energy storage by increasing glucose uptake and suppressing lipolysis. This metabolic response to insulin is mediated by an extensive signalling network^1^, which elicits cellular responses that require changes in protein subcellular localisation.

Insulin binding to its receptor, INSR, alters protein subcellular distribution through several mechanisms, including regulating protein-protein interactions via protein phosphorylation, lipid messenger generation (e.g., PI(3,4,5)P_3_), and direct effects on membrane trafficking. For example, recruitment of insulin receptor substrate 1 (IRS-1) and phosphoinositide 3-kinase (PI3K) to the activated INSR is mediated by binding to phospho-tyrosine residues on INSR and IRS1, respectively, whilst insulin-stimulated phosphorylation of the tuberous sclerosis complex (TSC)1/2 complex promotes its dissociation from the lysosomes^2^. Insulin signalling also directly targets known regulators of membrane trafficking (e.g., TBC1D4) to deliver glucose transporters (i.e., GLUT4) to the plasma membrane (PM) and promote glucose uptake^3^. Endosomal cargoes including transferrin receptor (TfR)^4,5^, insulin-like growth factor 2 receptor (IGF2R)/cation-independent mannose 6-phosphate receptor (CI-M6PR)^6,7^, and sodium/hydrogen exchanger 6 (NHE6)^8^ also undergo insulin-stimulated trafficking to the PM, although the physiological significance of endosomal cargo redistribution to the cell surface remains unclear. Collectively, these examples highlight how multiple mechanisms governing protein relocalisation play an essential role in the cellular insulin response. Moreover, insulin-stimulated protein redistribution is impaired in states of insulin resistance (e.g., GLUT4 delivery to the PM^9^), exemplifying the importance of regulating protein localisation in metabolic health.

Unbiased approaches to study the acute adipocyte insulin response have, to-date, focussed solely on protein phosphorylation^1,10^, while studies into insulin-regulated protein movement have been limited to studies on single proteins or identifying translocation to/from a single organelle^8,11^. Here, we combined unbiased global (Localisation of Organelle Proteins by Isotope Tagging after Differential ultraCentrifugation (LOPIT-DC)^12^) and targeted plasma membrane profiling^13^ subcellular proteomics techniques to map the spatial proteome of 3T3-L1 adipocytes and identify changes in protein localisation in response to an acute insulin stimulus. These studies revealed >500 high-confidence insulin-responsive proteins. To demonstrate the utility of identifying proteins that move in response to insulin, we focussed on C3ORF18, which redistributed to the plasma membrane following insulin stimulation. Studies in adipocytes depleted of C3ORF18 revealed that C3ORF18 is a previously uncharacterised regulator of adipocyte insulin signalling. Overall, our study provides a key resource highlighting the dynamic adipocyte subcellular proteome and impetus for future mechanistic studies into how insulin regulates adipocyte function.

## Results

### Mapping the subcellular 3T3-L1 adipocyte proteome using LOPIT-DC

To map the 3T3-L1 adipocyte proteome, we modified the LOPIT-DC subcellular fractionation workflow^12^ to include an additional lipid droplet-enriched fraction (Fig. 1a). Mass spectrometry analysis of 9 fractions obtained by differential centrifugation and the lipid droplet fraction identified 4269 proteins in non-stimulated (basal) replicates (n = 3) and 4384 proteins in insulin-stimulated replicates (n = 3), with 4005 proteins identified in both conditions (Fig. 1b). Using a set of organelle marker proteins (total 531; Supplementary Table 1), we confirmed organelles were uniquely distributed across fractions (Fig. 1c, Extended Data Fig. 1a) and formed distinct clusters when visualised using linear dimensionality reduction (PCA) (Fig. 1d). In addition, the lipid droplet fraction was enriched for known lipid droplet-localised proteins including the perilipins (PLIN1, PLIN2, PLIN3, PLIN4), VPS13C, CIDEC, galectin-12, and ABHD5 (Extended Data Fig. 1b,c).

**Figure 1:**
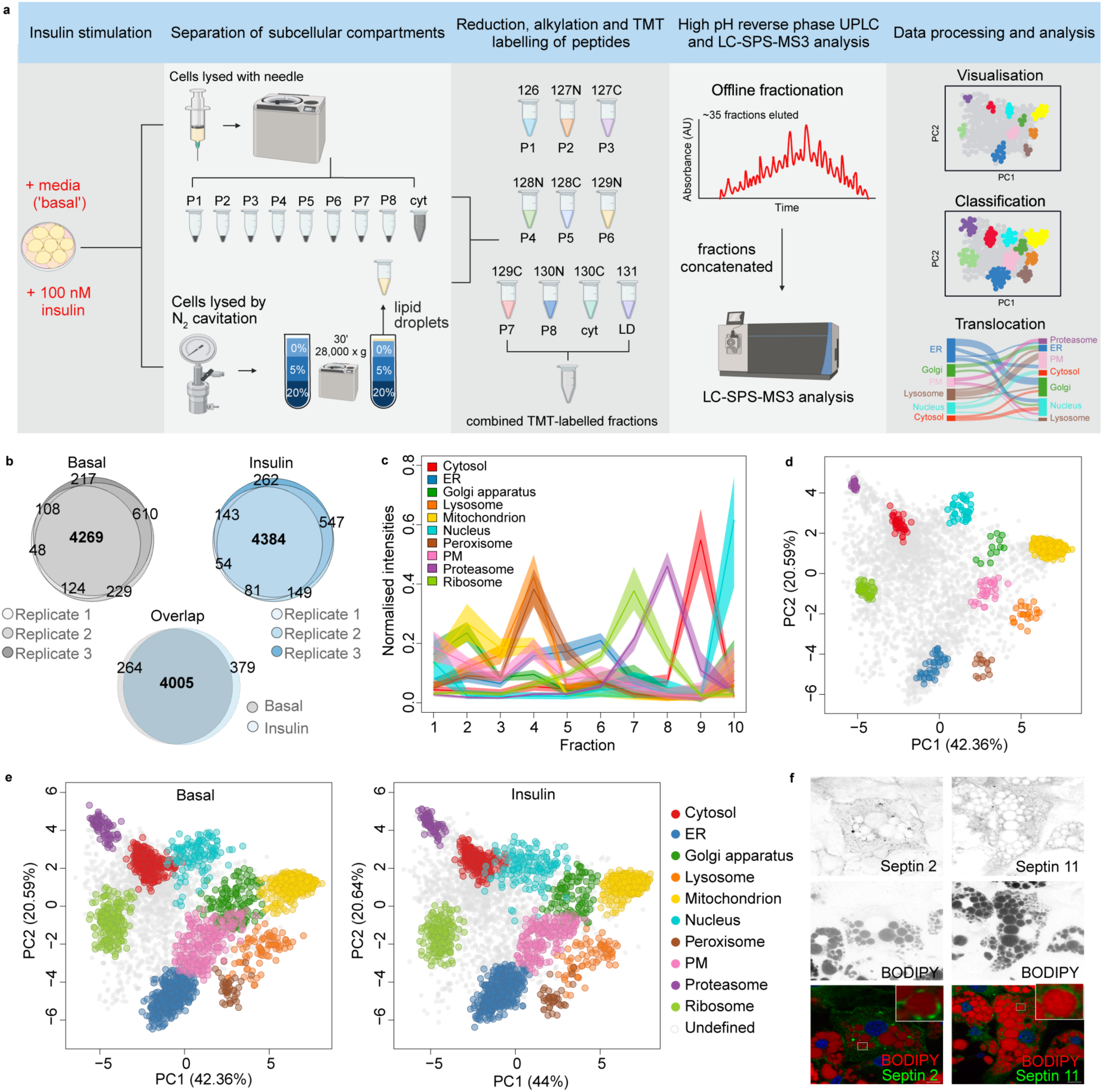
LOPIT-DC maps the spatial proteome of 3T3-L1 adipocytes under basal and insulin-stimulated conditions. a) Schematic of the experimental workflow used. b) Venn diagrams of overlap in proteins identified across three replicates of LOPIT-DC in basal and insulin-stimulated conditions. Data for each condition was concatenated to give 4005 proteins in common across both conditions. c) Profile of organelle marker proteins across all 10 fractions (basal replicate 1 data shown, all replicates in Extended Data Fig. 1a). d) Principal component analysis (PCA) projection of concatenated basal data (n = 3) showing distinct clustering of organelle marker proteins. Each point represents an individual protein. e) PCA projection of proteins allocated to each organelle under basal and insulin-stimulated conditions. f) Immunostaining of septin 2, septin 11 and BODIPY^TM^ 493/503 in 3T3-L1 adipocytes (scale bar = 10 μM).

Proteins were probabilistically assigned to a subcellular compartment based on their distribution across fractions using the machine learning algorithm BANDLE^14^. Accurate assignment requires organelles to have ≥10 marker proteins in all replicates with a consistent and unique distribution across fractions^14,15^. These criteria excluded the use of both lipid droplets and endosomes as organelles for protein allocation using BANDLE. Of the 4005 proteins, 1755 proteins in the basal dataset and 1614 in the insulin-stimulated data were assigned to a single subcellular location with high-confidence, in addition to the designated marker proteins (Fig. 1e; Supplementary Table 1). Of the allocated proteins, 1367 shared the same localisation under basal and insulin-stimulated conditions. The remaining ∼50% of proteins not allocated to an organelle were classified as ‘undefined’. A majority of these are likely multi-localised proteins found in more than one subcellular domain, as previously reported^8^. Protein localisations in basal and insulin-stimulated adipocytes can be viewed interactively at https://proteome.shinyapps.io/adipocyte2025/.

Gene ontology cellular compartment (GoCC) enrichment analysis of proteins allocated to specific subcellular compartments showed enrichment of expected terms (Supplementary Table 1). Comparing our protein organelle assignments to localisations in cultured human adipocytes^16^ (Simpson–Golabi–Behmel syndrome (SGBS) adipocytes; Supplementary Table 1) revealed high concordance, particularly for proteins allocated to organelles typically well resolved by centrifugation approaches (i.e., mitochondria, cytosol) (>80%) (Extended Data Fig. 1d). Organelles that are harder to resolve, such as those comprising the secretory pathway (endoplasmic reticulum (ER)-Golgi-PM), had lower concordance, likely reflecting dynamic protein traffic and multi-localisation within this pathway. Accordingly, a majority of proteins localised to the ER, Golgi or PM in 3T3-L1 adipocytes were assigned to organelles within the secretory pathway in SGBS adipocytes (Extended Data Fig. 1d).

Specific analysis of the lipid droplet fraction (fraction 10) revealed enrichment of all septin proteins identified (septin 2, 5, 6, 7, 8, 9, 10, 11) (Extended Data Fig. 1e) and these GTP– binding cytoskeletal proteins clustered near to lipid droplet marker proteins when visualised by PCA (Extended Data Fig. 1f). Septin 9 has been implicated in lipid droplet formation and localisation^17,18^ in non-adipocytes, whilst septins 7 and 11 are reported to regulate adipocyte lipid storage and metabolism^19–21^. Immunostaining of septin 2 and 11 in adipocytes revealed diffuse staining, but also distinct puncta close to lipid droplets (Fig. 1f). These data, in combination with their enrichment in the lipid droplet fraction, suggest septins localise to mature lipid droplets in adipocytes. Along with significant correlations between adipose tissue septin protein abundance and adipose-related clinical traits (Extended Data Fig. 1g)^22^, our data support the emerging role for this family of proteins in adipocyte biology. Overall, we provide a high resolution map of the 3T3-L1 adipocyte proteome which has high concordance to cultured human adipocytes and use these data to identify that septins localise to lipid droplets in mature adipocytes.

### Insulin stimulates widespread protein relocalisation

Having mapped the adipocyte proteome under basal and acute insulin-stimulated conditions, we next identified proteins with altered localisation in response to insulin. A Bayesian analysis framework was used to compare distribution profiles for proteins in unstimulated (basal) and insulin-stimulated adipocytes. This analysis identified 899 out of 4005 proteins as candidate movers (differential localisation probability (*diff. loc. prob.*) > 0), of which 502 were high-confidence (*diff. loc. prob.* = 1) (Fig. 2a; Supplementary Table 1). Known insulin-responsive proteins including GLUT4^44^ and LNPEP/IRAP^45,46^ were predicted to move with high-confidence (*diff. loc. prob.* = 1, Fig. 2b), validating our approach to detect insulin-responsive protein translocation.

**Figure 2:**
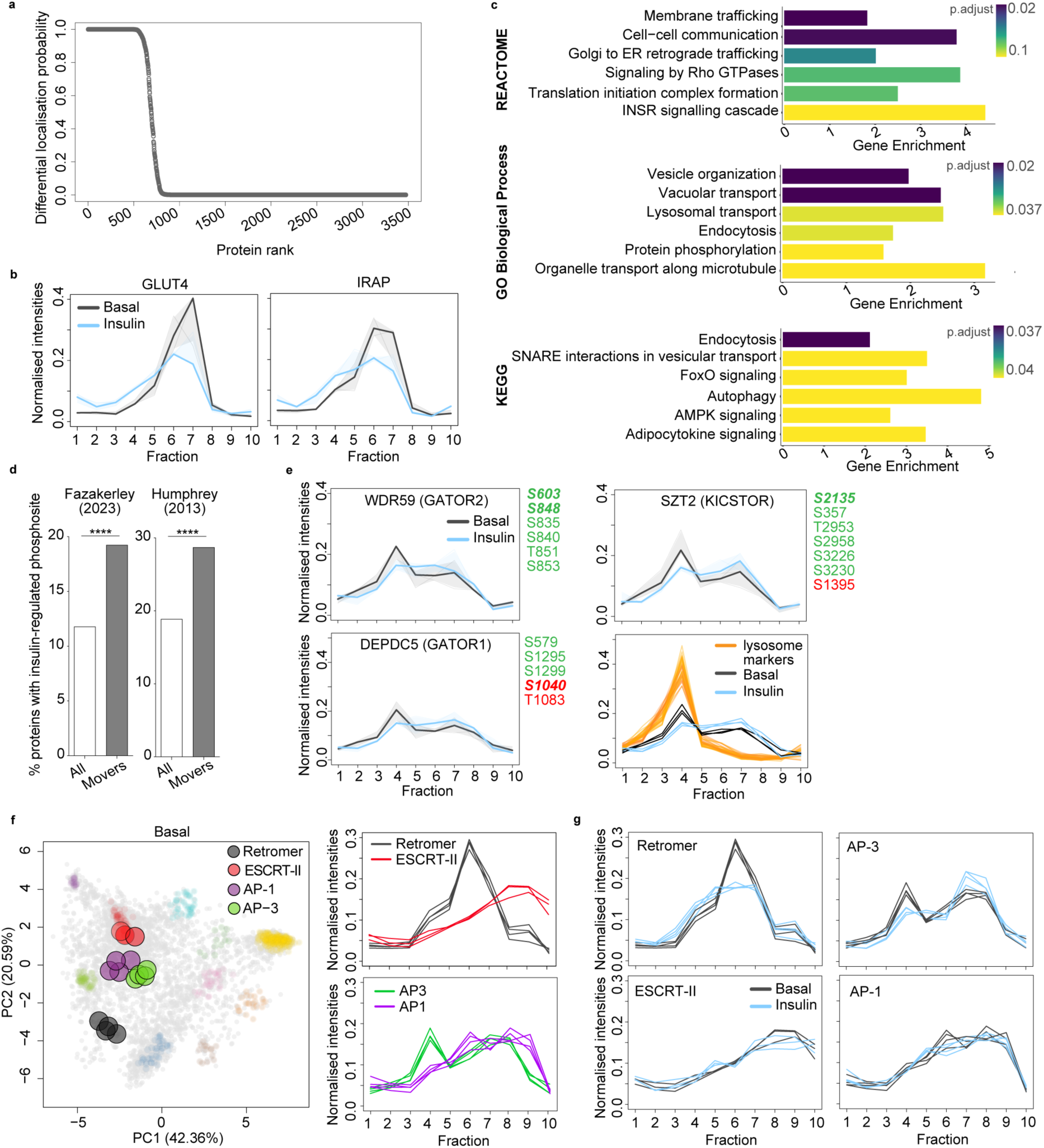
Insulin stimulates extensive changes in protein localisation, including proteins involved in cellular signalling and membrane trafficking. a) Differential localisation probability protein of all proteins (not including marker proteins) as determined using BANDLE. b) Distribution of GLUT4 and IRAP across fractions under basal (grey) and insulin-treated (blue) conditions (solid line = mean, shaded area 95% confidence interval (CI), n = 3 per condition). c) Gene set enrichment analysis of high confidence differentially localised proteins (*diff. loc. prob.* = 1) using Gene Ontology Biological Process, Reactome and KEGG databases (top 6 terms shown). d) Enrichment of proteins with insulin-regulated phosphorylation sites in 3T3-L1 adipocytes as reported by Fazakerley (2023)^10^ and Humphrey (2013)^1^ in all proteins vs. high-confidence differentially localised proteins (*diff. loc. prob.* = 1). Fisher’s exact test, Fazakerley *p* = 2.2 x 10^-11^, Humphrey *p* = 1.3 x 10^-13^. e) Distribution of DEPDC5 (GATOR1), WDR59 (GATOR2) and SZT2 (KICSTOR) subunits across fractions under basal (grey) and insulin-treated (blue) adipocytes (solid line = mean, shaded area 95% CI, n = 3 per condition), and insulin-regulated phosphorylation sites reported in Fazakerley (2023)^10^ and Humphrey (2013)^1^ (green = increased and red = decreased phosphorylation, regular font = found in either Fazakerley and Humphrey, bold italic = found in both Fazakerley and Humphrey). Bottom right panel shows WDR59, SZT2 and DEPDC5 profiles under basal (grey) and insulin-treated (blue) conditions with lysosomal protein markers in orange. f) PCA projection of concatenated basal LOPIT-DC data with retromer (black; VPS26A, VPS26B, VPS29, VPS35), ESCRT-II (red; VPS25, VPS36, SNF8), AP-1 (purple; AP1B1, AP1G1, AP1M1, AP1S1) and AP-3 (green; AP3B1, AP3D1, AP3M1, AP3S1) complex subunits highlighted (left panel), and mean distribution of subunits across fractions under basal conditions (right panel). g) Mean distribution of retromer, ESCRT-II, AP1 and AP3 subunits across fractions under basal (grey) and insulin-treated (blue) conditions (n = 3 per condition).

Differentially localised proteins were enriched for signalling terms (Fig. 2c), including insulin signalling, and for proteins with insulin-regulated phosphosites (Fig. 2d, Supplementary Table 1). The known insulin-regulated phosphoproteins IRS-1 and 2 were predicted to move with high confidence (*diff. loc. prob.* = 1, Extended Data Fig. 2A), whilst several protein kinases, including MARK2, SIK2, and mTOR, relocalised and were phosphorylated in response to insulin (*diff. loc. prob.* = 1, 1, 0.14, Extended Data Fig. 2b). In addition, several regulators of mTOR complex 1 (mTORC1) activity were also differentially localised and targeted by insulin signalling. These included the well-characterised target of insulin signalling, TSC1/2 (*diff. loc. prob.* = 1, Extended Data Fig. 2c), as well as members of the GATOR1 (DEPDC5), GATOR2 (WDR59) and KICSTOR (SZT2) complexes which are typically associated with the amino acid sensing branch of mTORC1 regulation (*diff. loc. prob.* = 0.99, 0.1, 0.4 respectively; Fig. 2e). Notably, DEPDC5, WDR59 and SZT2 exhibited a profile shift consistent with dissociation from lysosomes in response to insulin (Fig. 2e). Together, these data suggest that protein phosphorylation and redistribution work in concert to mediate adipocyte insulin signalling, and highlight the utility of assessing both post-translational modification and protein localisation in studying mechanisms of cell signalling.

Membrane trafficking terms were also over-represented in differentially localised proteins (Fig. 2c, Supplementary Table 1). These included proteins implicated in GLUT4^11,47,48^ (e.g., RALA, STX16, STX6, CDC42, PICALM, VTI1B, SCAMP1) and endosomal traffic (e.g., SNX12, VPS26A, SNX27, STX12) (Supplementary Table 1). We therefore assessed the localisation of cytosolic protein complexes that regulate cargo sorting through this system (e.g., retromer, ESCRT, AP-1, AP-2, AP-3, COPI, COPII, HOPS). These complexes had distinct distributions, and subunits within each complex clustered tightly when visualised by PCA (Fig. 2f, Extended Data Fig. 2c). Specific complexes had particularly striking relocalisation in response to insulin, including retromer (VPS26A/B-VPS29-VPS35, *mean subunit diff. loc. prob.* = 0.31; Fig. 2g) and AP-3 (*mean subunit diff. loc. prob.* = 0.77, Fig. 2g). However, others such as ESCRT-II and AP-1 did not alter their subcellular distribution (*mean subunit diff. loc. prob.* = 0, Fig. 2g). Together, these data suggest insulin signalling targets discrete components of the endolysosomal system.

### The plasma membrane proteome undergoes dynamic remodelling in response to insulin

LOPIT-DC revealed the adipocyte PM proteome to be highly insulin responsive (Fig 3a, Extended Data Fig. 3a). Therefore, we used an additional proteomics approach^13^ to quantify insulin-regulated changes to the PM proteome (Fig. 3b). 342 proteins exhibited altered abundance at the PM following insulin stimulation (119 increased; 223 decreased; Fig. 3c, Supplementary Table 2). These data highlight a substantial remodelling of the adipocyte PM proteome in response to acute insulin stimulation. We identified known insulin-responsive PM proteins GLUT1^49^, GLUT4^50–52^, LNPEP/IRAP^45,46^, IGF2R/CI-M6PR^6,7^ and TfR^4^ which all increased in abundance at the PM, and INSR^53^, which decreased (Fig. 3c). Twenty-seven proteins that changed in abundance at the PM in response to insulin (p<0.05) were also predicted to differentially localise with high-confidence in our whole cell subcellular proteomics data (*diff. loc. prob*. = 1) (Extended Data Fig. 3b, Supplementary Table 2). Within these, C3ORF18, an uncharacterised protein of unknown function, had one of the greatest (1.8-fold) increases in abundance at the PM in response to insulin (Supplementary Table 2). Accordingly, in our LOPIT-DC data, C3ORF18 also relocalised in response to insulin (*diff. loc. prob.* = 1, Fig. 3d), with greater enrichment in fractions containing PM marker proteins after insulin stimulation (Fig. 3e).

**Figure 3:**
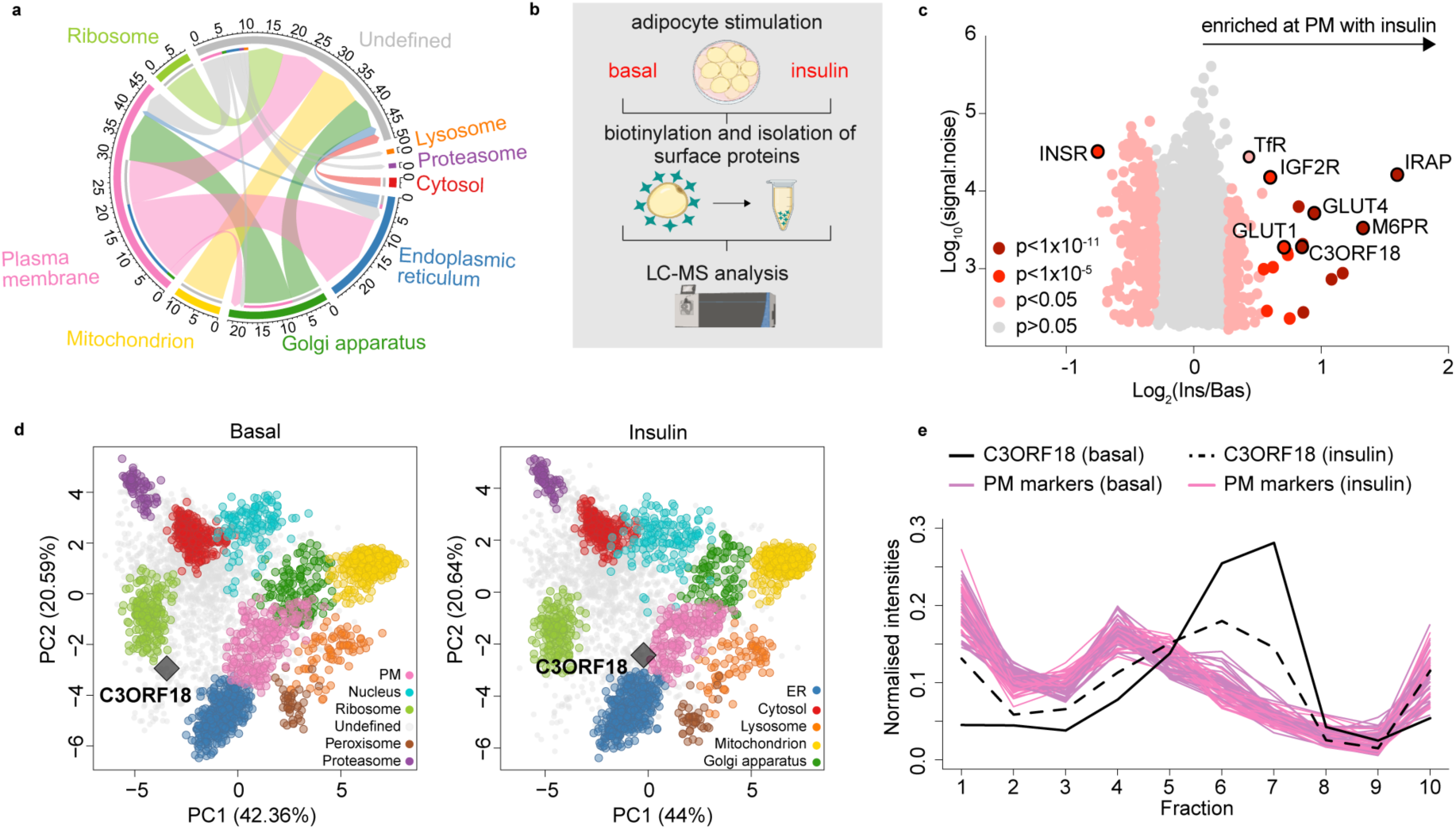
Quantitative plasma membrane proteomics reveals the extent of plasma membrane remodelling in response to insulin. a) Chord plot visualising movement of proteins predicted to relocalise with high-confidence (*diff. loc. prob*. = 1) assigned to an organelle in at least 1 condition. b) Schematic of plasma membrane proteomics workflow. c) Scatter plot of protein abundance at the 3T3-L1 adipocyte PM following insulin stimulation. *P*-values were calculated using Significance A values and corrected for multiple hypothesis testing using the Benjamini-Hochberg method. d) PCA projection of LOPIT-DC adipocyte subcellular map under basal (left) and insulin-stimulated (right) conditions highlighting C3ORF18 localisation (grey diamond). e) Distribution of PM marker proteins (pink = basal, purple = insulin) and C3ORF18 in LOPIT-DC data under basal (solid black line) and insulin-stimulated (dashed black line) conditions. Mean profile of each protein shown, n = 3 per condition.

### C3ORF18 is a novel insulin-responsive protein

The type III membrane protein C3ORF18 captured our attention as it underwent insulin-stimulated redistribution in both our whole-cell spatial proteomics and PM proteomic profiling, and was upregulated during adipocyte differentiation (Fig. 4a). We noted that despite a predicted molecular weight of ∼17 kDa, we detected an anti-C3ORF18 immunoreactive band at ∼40 kDa following SDS-PAGE and immunoblot. We validated this ∼40 kDa band as C3ORF18 by 1) observing the same apparent molecular weight with over-expressed epitope-tagged C3ORF18 (Extended Data Fig. 4a); 2) showing that siRNA-mediated depletion of C3ORF18 led to loss of this ∼40 kDa band (Extended Data Fig. 4b); and 3) demonstrating this ∼40 kDa band was insulin responsive in the PM fraction (Extended Data Fig. 4c).

**Figure 4:**
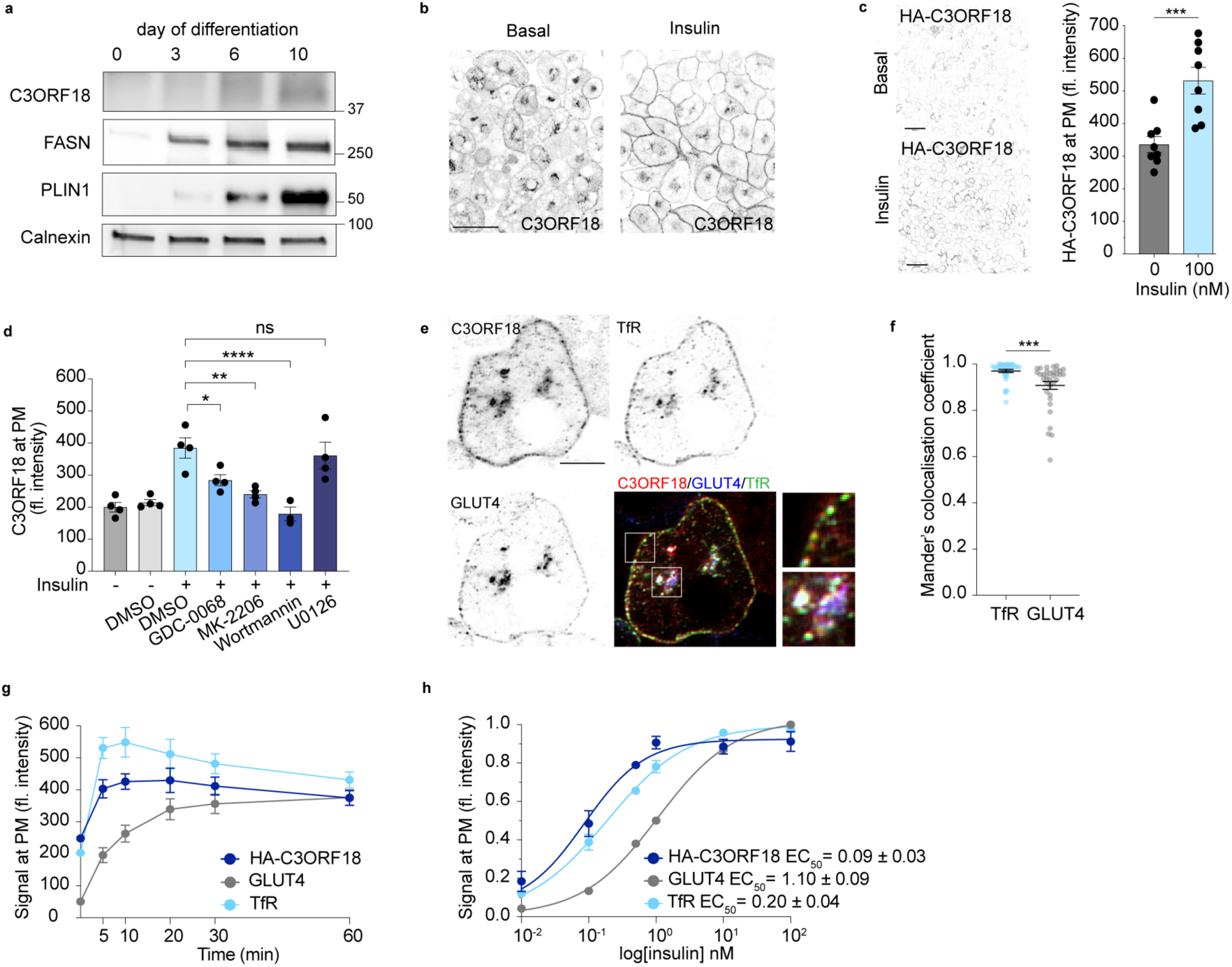
C3ORF18 is a novel insulin-responsive protein. a) Western blot analysis of C3ORF18 expression over differentiation in 3T3-L1 adipocytes. Representative blot, n = 2. b) Immunostaining of C3ORF18 in permeabilised 3T3-L1 adipocytes (scale bar = 50 μM) under basal and insulin-stimulated (100 nM, 30 min) conditions. c) Anti-HA surface immunostaining (left) and relative fluorescence intensity (right) in basal and insulin-stimulated (100 nM, 30 min) non-permeabilised 3T3-L1 adipocytes (n = 8 biological replicates, scale bar = 100 μM). d) Relative fluorescence intensity of surface HA-C3ORF18 in 3T3-L1 adipocytes treated with AKT-PI3K (GDC-0068, MK-2206, Wortmannin) or MAPK (U0126) signalling inhibitors prior to insulin stimulation (100 nM, 30 min, n = 3-4 biological replicates). e) Representative image of C3ORF18, TfR and GLUT4 immunostaining in 3T3-L1 adipocytes (scale bar = 10 μM) and f) quantification of colocalisation C3ORF18 with TfR and GLUT4 in (e) using Manders colocalisation coefficient in ImageJ, each data point represents an individual cell. g) Relative fluorescence intensity of surface HA-C3ORF18, GLUT4 and TfR in response to 100 nM insulin for 0-60 min in adipocytes (n = 7 biological replicates). h) Relative fluorescence intensity of surface HA-C3ORF18, GLUT4 and TfR in adipocytes following 30 min stimulation with increasing doses of insulin (0-100 nM, n = 7 biological replicates). All data represented as mean ± SEM (c, d, f-h). n.s. non-significant; \**p* < 0.05; \*\**p* < 0.01; \*\*\**p* < 0.01; \*\*\*\**p* < 0.0001 by paired two-tailed Student’s *t*-test (c, f) or one-way ANOVA with Šidák’s multiple comparisons test (d).

To validate the insulin-responsive relocalisation observed in our proteomics (Fig. 3d-f) and Western blot data (Extended Data Fig. 4c), we immunostained endogenous C3ORF18 in adipocytes under basal and insulin-stimulated conditions. This revealed C3ORF18 localised to both the perinuclear region and cell periphery, with increased peripheral staining in response to insulin (Fig. 4b). We observed similar localisation in adipocytes over-expressing HA-C3ORF18 (Extended Data Fig. 4d). To more accurately quantify the insulin-responsive translocation of C3ORF18 to the PM, we used N-terminally HA-tagged C3ORF18 (HA-C3ORF18) expressing cell lines, where the HA-epitope is lumenal/extracellular, to specifically label PM-localised C3ORF18. Surface staining in non-permeabilised cells confirmed insulin-responsive translocation of HA-C3ORF18 to the PM in 3T3-L1 adipocytes (Fig. 4c), SGBS adipocytes and L6 myotubes and myoblasts (Extended Fig. 4e). Small molecule inhibition of PI3K (Wortmannin) and AKT (GDC-0068, MK-2206), but not MAPK (U0126), impaired insulin-stimulated HA-C3ORF18 PM translocation (Fig. 4d, Extended Data Fig. 4f), demonstrating C3ORF18 relocalisation is dependent on the AKT-PI3K branch of insulin signalling, as previously described for both GLUT4 and TfR^54,55^ (Extended Data Fig. 4g,h).

Since the profile of C3ORF18 was strikingly similar to GLUT4 in our subcellular proteomics data (Extended Data Fig. 4i), we next assessed whether C3ORF18 localised to the specialised GLUT4 compartment or another insulin-responsive endosomal compartment with TfR. C3ORF18, GLUT4 and TfR largely colocalised within the perinuclear region (Fig. 4e).

Quantitative analysis revealed greater colocalisation of C3ORF18 with TfR than GLUT4 (Fig. 4f), best exemplified when assessing C3ORF18 and TfR immunostaining towards the cell periphery under basal conditions (Fig. 4e). Consistent with these co-localisation data, the dose response and kinetics of C3ORF18 translocation to the PM were more closely aligned with TfR than those of GLUT4 (Fig. 4g,h). For example, the t_1/2_ for C3ORF18 and TfR was <1 min and 1 min, respectively, but 6.2 min for GLUT4. Collectively these data suggest C3ORF18 redistributes to the plasma membrane from an insulin-responsive endosomal compartment.

### C3ORF18 is required for maximal insulin signalling

Having validated the insulin-responsive redistribution of C3ORF18 to the PM, we next determined whether this protein played a functional role in the adipocyte insulin response. SiRNA-mediated depletion of *C3ORF18* reduced insulin-stimulated glucose transport in human and mouse adipocytes (Fig. 5a,b Extended Data Fig. 5a,b), and, consistent with this, impaired insulin-stimulated GLUT4 translocation to the PM (Fig. 5c). Notably, insulin-stimulated TfR translocation was also impaired (Fig. 5d), suggesting that loss of C3ORF18 suppressed the insulin response more generally and it was not specific to GLUT4. Indeed, insulin signalling to AKT was decreased in C3ORF18–depleted adipocytes (Fig. 5e, Extended Data Fig. 5c), suggesting that loss of C3ORF18 limits the l adipocyte insulin response via impaired proximal insulin signalling. To determine clinical relevance of our observations we assessed the correlation between adipose tissue *C3ORF18* expression and clinical metabolic traits. *C3ORF18* expression was lower in those with higher circulating blood glucose and HbA1c (Fig. 5f), and increased with weight loss (Extended Data Fig. 5d). These data are consistent with C3ORF18 improving adipose insulin sensitivity.

**Figure 5:**
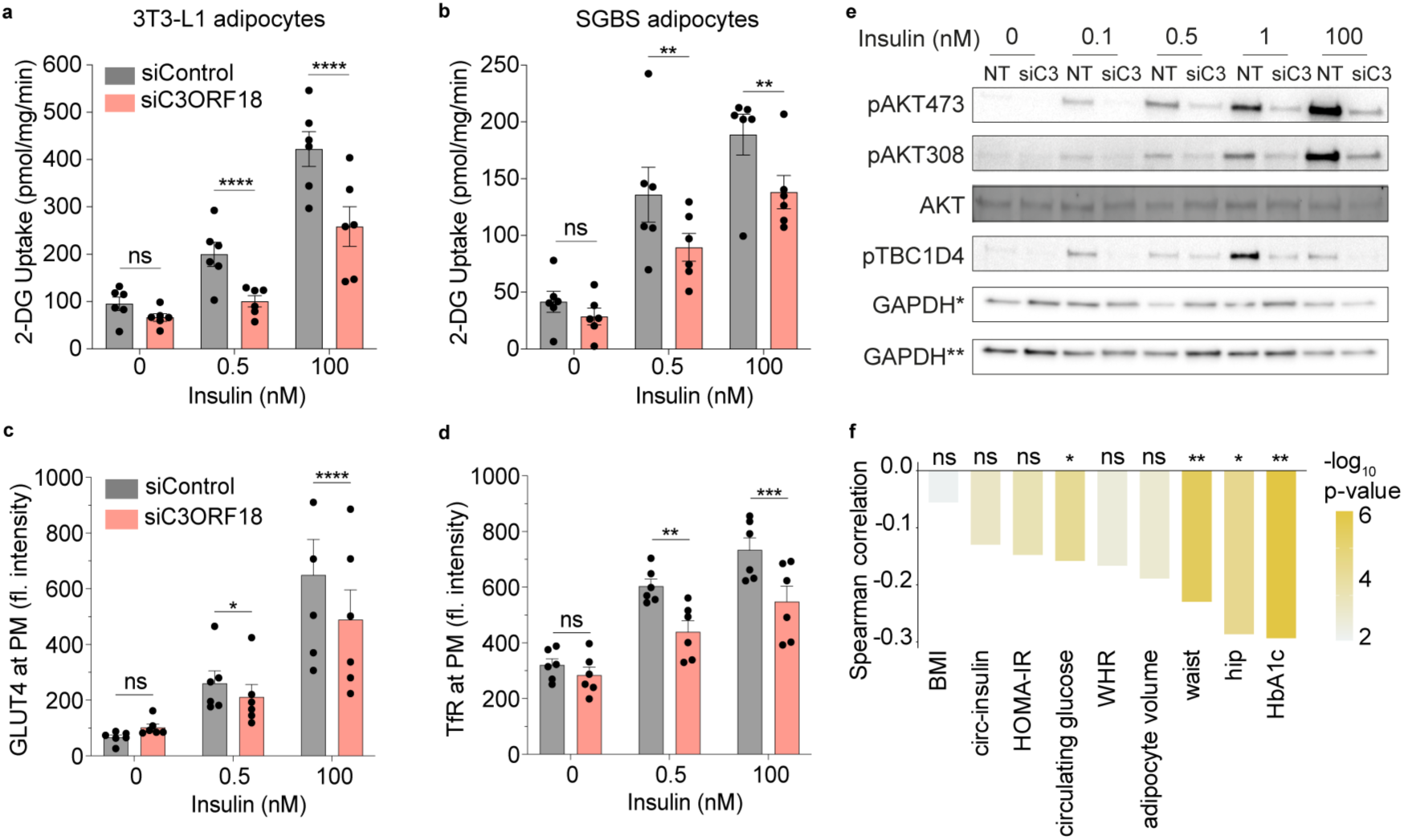
C3ORF18 is required for maximal insulin signalling. 2-deoxyglucose (2-DG) uptake in 3T3-L1 (a) and SGBS (b) adipocytes following C3ORF18 depletion (n = 6 biological replicates). Relative fluorescence intensity of surface GLUT4 (c) and TfR (d) in 3T3-L1 adipocytes stimulated with 0.5 or 100 nM insulin for 30 min (n = 6 biological replicates). e) Representative western blot of phosphorylated AKT (Thr308 and Ser473) and TBC1D4 (Thr642) in 3T3-L1 adipocytes following C3ORF18 depletion, stimulated with 0, 0.1, 0.5, 1 or 100 nM insulin for 30 min (n = 3-6 biological replicates). GAPDH* loading control for total AKT and pAKT308, GAPDH** loading control for pAKT473 and pTBC1D4, NT is non-targetting siControl. f) Spearman’s correlation of *C3ORF18* mRNA expression in omental adipose with metabolic clinical features. Figure generated from adiposetissue.org^22^ using data from ^23–43^ (** pFDR < 0.01, * pFDR < 0.05). All other data represented as mean ± SEM (a-d). n.s. non-significant; \**p* < 0.05; \*\**p* < 0.01; \*\*\**p* < 0.01; \*\*\*\**p* < 0.0001 by two-way ANOVA with Sidak’s multiple comparisons test (a-d).

## Discussion

Here, we present a subcellular map of the 3T3-L1 adipocyte proteome under basal conditions and following acute insulin stimulation, to provide the first unbiased cell-wide analysis of insulin-stimulated protein relocalisation in adipocytes. Combined with a quantitative analysis of the plasma membrane, these data revealed extensive protein redistribution in response to insulin. We highlight the utility of this dynamic subcellular mapping to identify novel regulators of adipocyte insulin action by studying C3ORF18, which traffics to the plasma membrane in response to insulin and is required for maximal insulin signalling in adipocytes.

Previous work has focused on insulin-stimulated protein translocation events in individual organelles, such as GLUT4 storage vesicles^11,48^ or the plasma membrane^8^, but no studies to-date have taken the unbiased whole-cell approach used herein. Approximately ∼10% (502 proteins) of identified proteins exhibited high-confidence (*diff. loc. prob. = 1*) subcellular redistribution in response to insulin, with the PM proteome particularly dynamic. We identified a number of novel insulin responsive proteins worthy of functional follow up, including the uncharacterised protein C3ORF18. Insulin-stimulated the redistribution of C3ORF18 to the PM from a TfR-positive intracellular compartment, likely endosomes, in adipocytes (mouse and human) and muscle cells. Depletion of C3ORF18 impaired insulin-stimulated glucose transport and delivery of both TfR and GLUT4 to the plasma membrane in adipocytes. This appears to be driven by impaired insulin signalling to AKT, suggesting that C3ORF18 may play a role in promoting or maintaining proximal insulin signalling. Consistent with this, *C3ORF18* expression in omental adipose tissue was negatively correlated with circulating glucose and HbA1c. Further, the link between C3ORF18 and adipocyte biology is supported by human genetics where a common variant, rs1034405, is associated with increased waist-to-hip ratio adjusted for BMI^56^. However, the molecular mechanisms linking C3ORF18 to insulin signalling remain unclear, and further work will be needed to interrogate C3ORF18 biochemistry (i.e., why it has a higher apparent molecular weight by Western blot), how its localisation is regulated by PI3K/AKT signalling, and how C3ORF18 promotes the adipocyte response to insulin signalling. Nonetheless, this example demonstrates that changes in protein subcellular location can be used to identify functionally relevant proteins.

The large effect on the plasma membrane proteome is, to some extent, unsurprising, since insulin-regulated delivery of GLUT4 (and proteins that colocalise with GLUT4) to the plasma membrane is highly studied^3^. However, a large proportion of proteins that increase in abundance at the PM with insulin are not reported to localise to specialised GLUT4 storage vesicles^11,47,48^, suggesting that other vesicular carriers are insulin responsive. Indeed, endosomal cargoes including IGF2R/CI-M6PR^6,7^, TfR^4,5^, and C3ORF18 as reported here, also undergo insulin-responsive trafficking to the PM. However, the extent of the endosomal response to insulin, why this occurs and how this is regulated remains unclear. Our data suggest that there is redistribution of several endosomal cargoes to the PM in response to insulin, and that insulin signalling specifically regulates a subset of endomembrane trafficking complexes which directly bind cargoes (for example retromer and AP-3). These data are in line with the growing recognition of the role the endosomal network plays in metabolic regulation and pathologies^57^, and provide a foundation for studies into the spatial regulation of endolysosomal trafficking by insulin signalling.

The scale of protein redistribution we identified highlights the likely importance of protein localisation in the adipocyte insulin responses. Proteins predicted to relocalise in response to insulin were often also subject to insulin-regulated phosphorylation, suggesting that, in some cases, these two mechanisms of protein regulation may be linked. As an example, mTOR itself, as well as subunits of the KICSTOR and GATOR complexes implicated in regulation of mTORC1 activity at the lysosomes, were both phosphorylated and moved away from lysosomes in response to insulin. As these complexes are primarily implicated in amino acid sensing, this raises the possibility that insulin signalling directly regulates mTORC1 activity via phosphorylation-driven redistribution of GATOR/KICSTOR, as recently reported for GATOR2 by AMPK signalling^58^. In this way, the combination of dynamic protein localisation and phosphorylation data may reveal new mechanisms governing the cellular insulin response.

There are several limitations to our study. Although the 3T3-L1 adipocyte model has been used extensively to study adipocyte cell biology, these cells are derived from mice and so may not fully recapitulate protein localisation in human adipocytes. However, our comparison with localisations reported in human SGBS adipocytes indicated a high degree of similarity in protein localisation between mouse and human adipocytes. Further, follow up work on C3ORF18 was carried out in both mouse and human cells, where the same phenotype was observed. We employed strict thresholding of allocations to avoid over-allocation of multi-localised proteins to single organelle(s), and endosomes and the lipid droplet were removed as organelles for allocation, likely contributing to the lower number of proteins allocated to a single organelle in our study compared to others^16^. Importantly, detection of protein relocalisation was independent of protein allocation to an organelle, a key benefit of BANDLE. Indeed, a majority of insulin-responsive proteins were not allocated to a specific organelle, and were of ‘undefined’ localisation under both conditions. These are likely multi-localised proteins, where a change in protein distribution with insulin represents changes in the relative abundance of the protein between subcellular sites. Finally, we generated data from two distinct subcellular proteomics approaches. Since these data were not entirely overlapping, there are likely many additional plasma membrane translocating proteins identified in our plasma membrane proteomic analysis that are worthy of follow-up, but were not considered here if not present in both analyses.

In summary, we present a high resolution subcellular map of the adipocyte proteome and demonstrate the extent and importance of insulin-stimulated protein relocalisation in the adipocyte insulin response. These data are available through an <u>interactive online resource</u> for further mining by the research community. Finally, we demonstrate C3ORF18 is required for a maximal insulin response in adipocytes, illustrating the utility of studying acute protein relocalisation to identify novel regulators of insulin action.

## Methods

### Cell Culture

#### 3T3-L1 fibroblast culture and differentiation into adipocytes

3T3-L1 fibroblasts obtained from David James (University of Sydney), originally from Howard Green (Harvard Medical School, Boston, MA), were maintained in Dulbecco’s modified Eagle’s medium (DMEM; Sigma) supplemented with 10% FBS (Gibco) and 2 mM GlutaMAX (Gibco) at 37°C with 10% CO_2_. Cells were passaged at ∼60% confluence. Differentiation was induced at 100% confluence by the addition of 350 nM insulin, 0.5 mM 3-isobutyl-1-methylxanthine (IBMX), 250 nM dexamethasone and 400 nM biotin to the supplemented DMEM for 3 d, followed by 350 nM insulin for a further 3 d. Adipocytes were used 10-11 d following the initiation of differentiation.

#### SGBS preadipocyte culture and differentiation into adipocytes

SGBS adipocytes were provided by Professor Martin Wabitsch (Division of Pediatric Endocrinology and Diabetes, University Medical Center Ulm, Germany), and were cultured and differentiated according to the established protocol^59,60^. Briefly, SGBS fibroblasts were maintained in DMEM/F-12 (Sigma) supplemented with 10% FBS, 2 mM GlutaMAX, 33 μM biotin and 17 μM pantothenate. Differentiation was induced at 100% confluence by the addition of 0.01 mg/mL transferrin, 20 nM insulin, 100 nM cortisol, 0.2 nM T3, 25 nM dexamethasone, 250 μM IBMX, and 2 μM rosiglitazone to serum-free DMEM/F-12 for 4 d. Adipocytes were maintained in post differentiation media containing 0.01 mg/mL transferrin, 20 nM insulin, 100 nM cortisol and 0.2 nM T3 in serum-free DMEM/F-12 until use at day 12-14 after initiation of differentiation.

#### L6 myoblast culture and differentiation into myotubes

L6 myoblasts obtained from David James (University of Sydney) were maintained in AlphaMEM (Gibco) supplemented with 10% FBS and 2 mM GlutaMAX. Differentiation was induced at 80-90% confluence by culture in AlphaMEM supplemented with 2% horse serum (PAN-Biotech, P30-0702) and 2 mM GlutaMAX, and media changed every 2 days. Cells were used 5-7 days following onset of differentiation.

#### Generation of HA-C3ORF18 overexpressing stable cell lines

Empty vector or HA-C3ORF18-overexpressing 3T3-L1 and L6 cell lines were generated via retroviral transduction (pMMLV; VectorBuilder), as previously described^61^. Retrovirus was generated using the Plat-E packaging cell line^62^. HA-C3ORF18-overexpressing SGBS cells were generated via lentiviral transduction (pLV; VectorBuilder). Lentivirus was generated using the LentiX 293T packaging cell line (Takara Bio, 632180). Transduced cells were selected with 2 μg/mL puromycin and maintained and differentiated as described above. Puromycin was removed prior to cell differentiation.

#### HEK cell culture and transient HA-C3ORF18 overexpression

Wild type HEK-293 cells were cultured in high glucose DMEM supplemented with 10% FBS (Gibco) and 2 mM GlutaMAX (Gibco) at 37°C with 5% CO_2_. Cells were seeded at 3 x 10^5^ cells/well into a 24-well plate 24 h prior to transfection with 0.5 μg DNA delivered using Lipofectimine2000 (Invitrogen). Cells were harvested for immunoblotting 24 h following transfection as described below.

### LOPIT-DC

#### Subcellular fractionation

Differentiated 3T3-L1 adipocytes were cultured in low media volume as previously described for 48 h prior to the experiment (8 mL/10 cm dish, media changed every 24 h) to improve insulin sensitivity^63^. serum starved in basal DMEM (high glucose DMEM, GlutaMAX, 0.2% BSA) for 2 h and maintained in basal medium or stimulated with 100 nM insulin for 30 min. Four confluent 10 cm dishes were used per replicate per condition. Cells were washed twice with ice-cold PBS and harvested in homogenisation buffer (250 mM sucrose, 10 mM 4-(2-hydroxyethyl)-1-piperazineethanesulfonic acid (HEPES) pH 7.4, 2 mM EDTA, 2 mM magnesium acetate tetrahydrate) containing EDTA-free phosphatase and protease inhibitor cocktail (Pierce). Cells were homogenised by 10 passes through a 21-gauge needle followed by 10 passes through a 27-gauge needle prior to centrifugation at 100 x g for 5 min at 4°C to remove unlysed cells. Cells were fractionated by centrifugation in a benchtop Beckman Allegra X-15R centrifuge (fraction 1-3), benchtop Eppendorf 5415R centrifuge (fraction 4-5), and a Beckman Optima LE-80K ultracentrifuge with a TLA-55 rotor (fraction 6-8) at 4°C (Supplementary Table 1) and pellets were kept on ice.

Pellets were resuspended in solubilisation buffer (8 M urea, 50 nM HEPES pH 7.4, 0.2% SDS) and sonicated using a tip probe sonicator. Supernatant from the last centrifugation step was retained as the cytosolic fraction, precipitated with ten volumes of cold acetone overnight at -20°C and centrifuged at 16,000 x g for 20 min at 4°C. Supernatant was removed and the pellet air dried for 15 min before resuspension in solubilisation buffer.

#### Lipid droplet isolation

Lipid droplets were isolated using a modified version of the protocol described by Brasaemle and Wolins^64^. 3T3-L1 adipocytes were serum-starved in basal DMEM for 2 h, and treated with 100 nM insulin for 30 min as described above. Six confluent 10 cm plates of adipocytes were used per condition. Cells were washed twice with ice cold PBS and harvested in Tris-EDTA buffer (20 mM Tris-HCl (pH 7.4), 1mM EDTA, cOmplete protease inhibitor (Merck)), and disrupted by nitrogen cavitation (Parr Instruments). Cells were exposed to 41 Barr of nitrogen for 20 min, after which the pressure was slowly released and the cell homogenate collected dropwise. Homogenate was spun at 1000 x g for 10 min at 4°C to remove unlysed cells, the supernatant made up in 20% sucrose (w/v) in Tris-EDTA buffer and overlaid with 5% and 0% sucrose in Tris-EDTA buffer in a 4 mL ultracentrifuge tube (Beckman Coulter). Samples were centrifuged at 28,000 x g for 30 min at 4°C with no brake in a Beckman Coulter Optima LE-80K ultracentrifuge using a SW 40 Ti rotor (Beckman Coulter). Lipid droplets were harvested from the top of the gradient, washed once and resuspended in Tris-EDTA buffer. Lipid droplet associated proteins were precipitated using chloroform-methanol and the protein pellet air dried, before being resuspended in 100 μL solubilisation buffer and incorporated into the LOPIT-DC workflow.

#### Proteomic sample preparation

100 μg of protein from each fraction was reduced with final concentration 10 mM DTT at 37°C for 1 h, then alkylated with final concentration 40 mM chloroacetamide (CAA) for 2 h at RT. Protein was precipitated with acetone overnight at -20°C, and pellets resuspended in HEPES, pH 8.5. Samples were digested with sequencing-grade trypsin (Promega) with a final enzyme:protein ratio of 1:40 for 1 h at 37°C, followed by a second trypsinisation step at 1:20 overnight at 37°C. Peptides were quantified using a fluorometric peptide assay (Thermo Fisher Scientific) according to manufacturer’s instructions.

50 μg of each fraction was labelled using a 10-plex tandem mass tag (TMT) kit (Thermo Fisher Scientific, Supplementary Table 1). TMT tags were equilibrated to RT and resuspended in 82 μL of LC-MS grade acetonitrile (ACN). 100 μL of each fraction (at concentration 0.5 μg/μL) was added to 41 μL of the appropriate tag. Labelling was allowed to proceed for 2 h on a shaker at RT. Reactions were quenched by adding 5% hydroxylamine and incubating for a further 45 min on a shaker at RT. The 10 fractions of each sample were then pooled into 1 LoBind Eppendorf and lyophilised using a SpeedyVac.

Samples were desalted using a C18 SepPak cartridge (100 mg sorbant, Waters). Cartridges were equilibrated with 2 washes of 100% ACN, 1 wash of 0.05% acetic acid and 3 washes of 0.1% trifluoroacetic acid (TFA). Samples were resuspended in TFA and the pH adjusted to < 3 before loading on column, and washed 3 times with 0.1% TFA, twice with 0.05% acetic acid, eluted in 70% ACN and 0.05% acetic acid and lyophilised.

Samples were pre-fractionated offline using reverse phase UPLC in an Acquity UPLC System with diode array detector (Waters) using an Acquity UPLC bridged ethyl hybrid C18 column (2.1-mm ID × 150-mm; 1.7-µm particle size; Waters). Peptide fractions were collected over a 50 minute linear gradient, combined into 15 concatenated fractions, and lyophilised.

#### LC-MS3

MS analysis was performed using an Lumos Orbitrap mass spectrometer coupled to a Dionex Ultimate 3000 RSLC nanoUPLC system (Thermo Fisher Scientific, Waltham, MA, USA). Peptides were loaded onto a pre-column (Thermo Fisher Scientific PepMap 100 C18, 5 mm particle size, 100 °A pore size, 300 mm i.d. x 5 mm length) from the Ultimate 3000 auto-sampler with 0.1% formic acid (FA) for 3 min at a flow rate of 15 μL/min. After this period, the column valve was switched to allow elution of peptides from the pre-column onto the analytical column. Separation of peptides was performed by C18 reverse-phase chromatography at a flow rate of 300 nL/min using a Thermo Fisher Scientific reverse-phase nano Easy-spray column (Thermo Fisher Scientific PepMap C18, 2 mm particle size, 100 °A pore size, 75 μm i.d. x 50 cm length). Solvent A was water + 0.1% FA and solvent B was 80% ACN, 20% water + 0.1% FA. The linear gradient employed was 2-40% B in 93 min (total LC run time was 120 min including a high organic wash step and column re-equilibration).

The eluted peptides from the C18 column LC eluant were sprayed into the mass spectrometer by means of an Easy-Spray source (Thermo Fisher Scientific). All m/z values of eluting peptide ions were measured in an Orbitrap mass analyser, set at a resolution of 120,000 and were scanned between m/z 380-1500 Da. Data dependent MS/MS scans (Top Speed) were employed to automatically isolate and fragment precursor ions by collision-induced dissociation (CID; normalised collision energy (NCE): 35%) which were analysed in the linear ion trap. Singly charged ions and ions with unassigned charge states were excluded from being selected for MS/MS and a dynamic exclusion window of 70 s was employed. The top 10 most abundant fragment ions from each MS/MS event were then selected for a further stage of fragmentation by synchronous precursor selection (SPS) MS3 in the HCD high energy collision cell using HCD (NCE: 65%). The m/z values and relative abundances of each reporter ion and all fragments (mass range from 100-500 Da) in each MS3 step were measured in the Orbitrap analyser, which was set at a resolution of 50,000. This was performed in cycles of 10 MS3 events before the Lumos instrument reverted to scanning the m/z ratios of the intact peptide ions and the cycle continued.

#### MS spectra processing and peptide and protein identification

Raw LC-MS data files were processed with Proteome Discoverer v2.5 (Thermo Fisher Scientific) using the Mascot server 2.8.3 (Matrix Science). The MS data was run against a Swiss-Prot Mus musculus database (downloaded September 2019) and the common repository of adventitious proteins (cRAP, v1.0). Precursor and fragment mass tolerances were set to 10 ppm and 0.2 Da, respectively. Trypsin was set as the enzyme of choice and a maximum of 2 missed cleavages were allowed. Fixed modifications were set to carbamidomethyl(C) TMT6plex (N-term) and TMT6plex (K), and variable modifications were set to carbamyl(N-term), carbamyl(K), carbamyl(R), deamidation(N,Q), oxidation (M) and TMT6plex (S/T). Percolator version 3.05 was used to assess the false discovery rate (FDR) and only high-confidence peptides were retained. Quantification of the MS3 reporter ions was performed within the Proteome Discoverer workflow using the Most Confident Centroid method for peak integration and integration tolerance of 20 p.p.m. Reporter ion intensities were adjusted to the batch-specific TMT reporter ion isotope distributions prior to processing (Supplementary Table X). The mass spectrometry proteomics data have been deposited to the ProteomeXchange Consortium via the PRIDE partner repository with the dataset identifier PXD061017.

#### Spatial proteomics analysis

All proteomics data analysis was performed in R (v4.3.2) using the Bioconductor packages pRoloc (v1.42.0)^15^, bandle (v1.6)^14^, clusterProfiler (v4.10.1)^65,66^ and ggplot2 (v3.5.1).

Peptide spectrum match (PSM)-level quantitation output from Proteome Discoverer was imported into R and non-specific filtering was performed as previously described^12,67^ to retain high quality PSMs. PSMs were sum normalised and aggregated to protein level intensities using the ‘robust’ method within the pRoloc R package^68^. Replicates were concatenated to yield 4005 proteins quantified across all 6 replicates and allow the generation of a single subcellular map for each condition. The data was annotated with markers from previous subcellular proteomics experiments^12,69,70^. Additional marker proteins were identified through examining protein localisation in UniProt for proteins reported only to be localised to one organelle, and by mining the literature. Collectively, this gave a robust list of proteins which gave distinct clusters when projected onto PCA and t-SNE plots. Some markers did not cluster with the other corresponding organellar markers, which may be due to heterogeneity of protein localisation between cell types, or biological or technical variation between replicates. These markers were identified by visualising the PCA/t-SNE plots on the interactive interface of pRolocGUI and removed. The final marker list incorporated 531 proteins from 9 subcellular niches; cytosol, ER, Golgi apparatus, lysosomes, mitochondrion, nucleus, peroxisomes, PM, and ribosomes (Supplementary Table 1).

The Bayesian ANalysis of Differential Localisation Experiments (BANDLE) algorithm from the bandle package^71^ was used to assign proteins to subcellular niches and predict differential localisation with the following parameters; 30,000 MCMC iterations, 5,000 burn-in iterations, 4 chains, and 1/20 thinning. Each protein was assigned to a subcellular niche for which it had the highest mean allocation probability, as described in the bandle vignette on Bioconductor (https://bioconductor.org/packages/release/bioc/html/bandle.html)^72^. Allocations were thresholded to retain only high confidence allocations. Proteins were assigned to a subcellular niche if their allocation probability was >0.9 and their outlier probability was < median outlier probability for that organelle. Proteins were deemed differentially localised if differential localisation probability > 0. Proteins with differential localisation probability = 1 were considered ‘high confidence’ and all others as ‘candidates’.

#### Enrichment analysis

Gene ontology^66,73^ cellular compartment enrichment was performed in turn for proteins assigned to each organelle (not including marker proteins) using clusterProfiler^65,66^. Gene set enrichment analysis was performed using Gene ontology biological process and KEGG^74^ enrichment in clusterProfiler, and REACTOME^75^ enrichment analysis was performed using reactomePA (v1.46). In all analyses *p*-values were adjusted using the Benjamini-Hochberg procedure^76^.

#### Phosphoproteomics enrichment analysis

Proteins in Humphrey (2013)^1^ with phosphorylation sites with -/+ 2 fold change in response to insulin at any time point (0.25, 0.5, 1, 2, 5, 10, 20, 60 min) and proteins from Fazakerley (2023)^10^ with phosphorylation sites significantly up- or down-regulated in response to insulin were considered “insulin-regulated”. This list was converted to mouse uniprot IDs using UniProt align, with only reviewed entries taken. Enrichment of insulin-stimulated phosphorylation in the proteins predicted to differentially localise vs. not was determined using a Fisher’s exact test in R. Insulin-stimulated phosphorylation status of all proteins identified in LOPIT-DC in both phosphorylation data sets are shown in Supplementary Table 1.

### Plasma membrane proteomics

Plasma membrane proteomics was performed as previously described^13^ with minor modifications. Differentiated 3T3-L1 adipocytes were cultured in low media volume as described above for 48h prior to the experiment, serum starved in basal DMEM for 2 h and maintained in basal medium or stimulated with 100 nM insulin for 30 min. Three confluent 10 cm dishes were used per replicate per condition, and six biological replicates were performed. Cells were washed twice with ice-cold PBS containing MgCl_2_ and CaCl_2_ (Sigma) and all subsequent steps were performed on ice. An oxidation/biotinylation mix comprising 200 mM aminooxybiotin (Biotium), 3 mM sodium meta-periodate (Thermo Fisher Scientific) and 10 mM aniline (Sigma) in ice-cold PBS pH 6.7 was added and dishes were rocked on ice for 1 h at 4°C in the dark. The reaction was quenched by adding glycerol to a final concentration of 1 mM for 5 min. Cells were washed twice with PBS pH 7.4 and lysed in a lysis buffer comprising 1.6% Triton X-100, 10 mM Tris-HCl pH 7.6, 150 mM NaCl, 5 mM iodoacetamide (IAA) and cOmplete protease inhibitor (Merck) for 30 min on ice. Nuclei (pellet) and lipid (top phase) were removed by centrifugation at 4°C, 13,000 x g for 10 min, which was repeated three times. 5 mg of protein was incubated with high-affinity streptavidin agarose beads (Pierce, 20357, 10 μL beads/mg protein) at 4°C for 18 h on a rotor to capture biotinylated glycoproteins. Beads were washed extensively with lysis buffer and PBS/0.5% SDS, using Poly-Prep columns (BioRad) attached to a vacuum manifold. Captured proteins were reduced with PBS/0.5% SDS/100 mM DTT for 20 min at RT, washed extensively with UC buffer (6 M urea in 0.1 M Tris-HCl pH 7.6), alkylated with 50 mM IAA (Sigma) for 20 min at RT, and digested on-bead with trypsin (Promega) in 200 mM HEPES pH 8.5 for 3 h at 37°C. Tryptic peptides were collected and stored at -80°C until TMT labelling.

#### Peptide labelling with tandem mass tags

TMT reagents (TMTpro 16-plex, 0.8 mg, Thermo Fisher Scientific) were dissolved in 43 μL anhydrous ACN and 10 μL added to peptide at a final ACN concentration of 30% (v/v). Samples were labelled as in Supplementary Table 2, incubated at RT for 1 h, and reaction quenched with hydroxylamine to a final concentration of 0.5% (v/v). TMT-labelled samples were combined at a 1:1:1:1:1:1:1:1:1:1:1:1 ratio. The sample was vacuum-centrifuged to near dryness and subjected to C18-based solid phase extraction (Sep-Pak, Waters). An unfractionated ‘single shot’ was analysed initially to ensure similar peptide loading across each TMT channel, thus avoiding the need for excessive electronic normalization. As all normalization factors were >0.5 and <2, data for each single-shot experiment was analysed with data for the corresponding fractions to increase the overall number of peptides quantified. Normalization is discussed in ‘data analysis’ and high pH reversed-phase (HpRP) fractionation is discussed below.

#### Offline HpRP fractionation

TMT-labelled tryptic peptides were subjected to HpRP fractionation using an Ultimate 3000 RSLC UHPLC system (Thermo Fisher Scientific) equipped with a 2.1 mm internal diameter (ID) x 25 cm long, 1.7 μLm particle Kinetix Evo C18 column (Phenomenex). Mobile phase consisted of A: 3% ACN, B: 100 % ACN and C: 200 mM ammonium formate pH 10. Isocratic conditions were 90% A/10% C, and C was maintained at 10% throughout the gradient elution. Separations were conducted at 45°C. Samples were loaded at 200 μL/min for 5 min. The flow rate was then increased to 400 μL/min over 5 min, after which the gradient elution proceeded as follows: 0-19% B over 10 min, 19-34% B over 14.25 min, 34-50% B over 8.75 min, followed by a 10 min wash at 90% B. UV absorbance was monitored at 280 nm and 15 s fractions were collected into 96 well microplates using the integrated fraction collector. Adjacent columns of fractions were combined, resulting in six combined fractions. Wells were excluded prior to the start or after the cessation of elution of peptide-rich fractions, as identified from the UV trace. Fractions were dried in a vacuum centrifuge and resuspended in 10 μL MS solvent (4% ACN/5% FA) prior to LC-MS3.

#### LC-MS3

Labelled samples were analysed using an Orbitrap Fusion Lumos (Thermo Fisher Scientific). An Ultimate 3000 RSLC UHPLC machine equipped with a 300 μm internal diameter × 5 mm Acclaim PepMap μ-Precolumn (Thermo Fisher Scientific) and a 75 μm internal dimeter × 50 cm 2.1 μm particle Acclaim PepMap RSLC analytical column were used. The loading solvent was 0.1% FA. The analytical solvent consisted of 0.1% FA (A) and 80% ACN + 0.1% FA (B). All separations were carried out at 40°C. Samples were loaded at 5 μL/min for 5 min in loading solvent. For the single-shot sample, the analytical gradient consisted of 3-7% B over 3 min, 7-37% B over 173 min, followed by a 4 min wash at 95% B and equilibration at 3% B for 15 min. For fractionated samples, the analytical gradient consisted of 3-7% B over 4 min, 7-37% B over 116 min, followed by a 4 min wash at 95% B and equilibration at 3% B for 15 min. Each analysis used a MultiNotch MS3-based TMT method ^77,78^. The following settings were used: MS1: 380-1500 Th, 120,000 resolution, 2×10^5^ automatic gain control (AGC) target, 50 ms maximum injection time. MS2: quadrupole isolation at an isolation width of mass-to-charge ratio (m/z) 0.7, collision-induced dissociation fragmentation (normalized collision energy (NCE) 34) with ion trap scanning in turbo mode from m/z 120, 1.5×10^4^ AGC target, 120 ms maximum injection time. MS3: in Synchronous Precursor Selection mode, the top 10 MS2 ions were selected for HCD fragmentation (NCE 45) and scanned in the Orbitrap at 60,000 resolution with an AGC target of 1×105 and a maximum accumulation time of 150 ms. Ions were not accumulated for all parallelisable time. The entire MS/MS/MS cycle had a target time of 3 s. Dynamic exclusion was set to ± 10 ppm for 70 s. MS2 fragmentation was triggered on precursors 5×10^3^ counts and above. The mass spectrometry proteomics data have been deposited to the ProteomeXchange Consortium via the PRIDE partner repository with the dataset identifier PXD061616.

#### Data analysis

Mass spectra were processed using a Sequest-based software pipeline for quantitative proteomics, ‘MassPike’, through a collaborative arrangement with Professor Steven Gygi’s laboratory at Harvard Medical School. MS spectra were converted to mzXML using an extractor built upon Thermo Fisher’s RAW File Reader library (version 4.0.26). In this extractor, the standard mzxml format has been augmented with additional custom fields that are specific to ion trap and Orbitrap mass spectrometry and essential for TMT quantitation. These additional fields include ion injection times for each scan, Fourier Transform-derived baseline and noise values calculated for every Orbitrap scan, isolation widths for each scan type, scan event numbers, and elapsed scan times. This software is a component of the MassPike software platform and is licensed by Harvard Medical School.

A combined database was constructed from the mouse Uniprot database (2019) and common contaminants such as porcine trypsin and endoproteinase LysC. The combined database was concatenated with a reverse database composed of all protein sequences in reversed order. Searches were performed using a 20-ppm precursor ion tolerance. Fragment ion tolerance was set to 1.0 Th. TMT tags on lysine residues and peptide N termini (304.2071 Da) and carbamidomethylation of cysteine residues (57.02146 Da) were set as static modifications, while oxidation of methionine residues (15.99492 Da) was set as a variable modification.

To control the fraction of erroneous protein identifications, a target-decoy strategy was employed^79^. Peptide spectral matches (PSMs) were filtered to an initial peptide-level FDR of 1% with subsequent filtering to attain a final protein-level FDR of 1%. PSM filtering was performed using a linear discriminant analysis, as described previously^79^. This distinguishes correct from incorrect peptide IDs in a manner analogous to the widely used Percolator algorithm (https://noble.gs.washington.edu/proj/percolator/), through employing a distinct machine learning algorithm. The following parameters were considered: XCorr, ΔCn, missed cleavages, peptide length, charge state, and precursor mass accuracy. Protein assembly was guided by principles of parsimony to produce the smallest set of proteins necessary to account for all observed peptides (algorithm described in^79^).

Proteins were quantified by summing TMT reporter ion counts across all matching peptide-spectral matches using ‘MassPike’, as described previously^78^. Briefly, a 0.003 Th window around the theoretical m/z of each reporter ion was scanned for ions and the maximum intensity nearest to the theoretical m/z was used. The primary determinant of quantitation quality is the number of TMT reporter ions detected in each MS3 spectrum, which is directly proportional to the signal-to-noise (S:N) ratio observed for each ion. An isolation specificity filter with a cut-off of 50% was additionally employed to minimize peptide co-isolation^78^.

Peptide-spectral matches with poor quality MS3 spectra (a combined S:N ratio of less than 250 across all TMT reporter ions) or no MS3 spectra at all were excluded from quantitation. Peptides meeting the stated criteria for reliable quantitation were then summed by parent protein, in effect weighting the contributions of individual peptides to the total protein signal based on their individual TMT reporter ion yields. Protein quantitation values were exported for further analysis in Excel.

For protein quantitation, reverse and contaminant proteins were removed, then each reporter ion channel was summed across all quantified proteins and normalised assuming equal protein loading across all channels. For further analysis and display in Figures, fractional TMT signals were used (i.e. reporting the fraction of maximal signal observed for each protein in each TMT channel, rather than the absolute normalised signal intensity). This effectively corrected for differences in the numbers of peptides observed per protein. *P* values were calculated using the method of significance A and corrected for multiple hypothesis testing in Perseus version 1.5.2.20.4^80^.

#### Immunoblot analysis of isolated PM fraction

Following incubation with high-affinity streptavidin agarose beads as described above, beads were washed with lysis buffer, PBS/0.5% SDS and UC buffer, and incubated with 4x Laemmeli Sample buffer (Bio-Rad) and tris(2-carboxyethyl)phosphine powder (TCEP; Thermo Fisher Scientific) at 95°C to elute proteins for immunoblot analysis.

### Immunoblotting

Adipocytes were cultured in reduced media volumes as previously described for 48 h prior to the experiment (250 μl media per well of 24-well plate, media changed every 24 h) to improve insulin sensitivity^63^. Cells were washed twice in ice-cold PBS and lysed in RIPA lysis buffer (50 nM Tris pH 7.5, 150 mM NaCl, 1 mM EDTA, 1% Triton, 0.5% Na Deoxycholate, 0.1% SDS, 1% glycerol) containing EDTA-free Phosphatase and Protease Inhibitor Cocktail (Pierce). Cell lysates were sonicated and centrifuged at 16,000 x g, 20 min, 4°C to remove lipid (top layer) and unlysed cells/debris (pellet). Protein concentration of the infranatant was quantified using a BCA Protein Assay Kit (Pierce) according to manufacturer’s instructions. Lysates were denatured by mixing with 4x Laemmeli Sample buffer, reduced with TCEP and heated at 65°C for 10 min. Protein was resolved by SDS-PAGE using 4-20% (or 10% for C3ORF18) Mini-PROTEAN TGX Stain-Free Gels (Bio-Rad) and transferred to a nitrocellulose membrane using the Trans-Blot Turbo Transfer System (Bio-Rad). Membranes were blocked in 5% (w/v) dried skimmed milk (Marvel) in tris-buffered saline for 1 h at RT, followed by overnight incubation at 4°C with the appropriate primary antibody (Supplementary Table 3). Membranes were incubated with the appropriate horseradish peroxidase (HRP) or fluorophore-conjugated secondary antibodies (Supplementary Table 3) diluted 1:5000 in 5% (w/v) dried skimmed milk for 1 h at RT. Signals were detected using enhanced chemiluminescence (ECL; Thermo Fisher Scientific) or in the appropriate fluorescence channel on a ChemiDoc MP Imaging System (Bio-Rad). Band intensities were quantified using ImageLab (v6.1, Bio-Rad). Data in Extended Fig. 5c was min-max normalised to the basal and 100 nM responses within replicates to account for varying signal total intensity between replicates.

### siRNA knockdown in differentiated adipocytes

siRNA knockdown in 3T3-L1 adipocytes was performed as previously described^81,82^. Briefly, siRNA targeting mouse C3ORF18 (6430571L13Rik, siGENOME SMARTpool #235599; Dharmacon Horizon) at final concentration 50 nM was delivered to differentiated adipocytes (day 6 following differentiation) using TransIT-X2 (Mirus Bio). Cells were fed with fresh media 24 h following transfection, and transfected again 96 h after the initial transfection as previously described^82^. Functional assays were performed 96 h following the second transfection.

SGBS adipocytes were transfected as described above, with minor modifications. Forward transfection with Lipofectamine RNAiMax (Thermo Fisher Scientific) was used to deliver siGENOME SMART pool #51161 (human C3ORF18) at final concentration 50 nM on day 10 and day 14 after differentiation. Functional assays were performed 96 h following the second transfection.

### qPCR

RNA extractions were performed using the RNeasy Mini kit (Qiagen 1152 #74104). Concentrations of RNA samples were quantified using NanoDrop. cDNA synthesis from 500 ng RNA was performed using the GoScript Reverse Transcriptase kit (Promega #A2801). Real-time (RT)-polymerase chain reaction (PCR) was performed using TaqMan or SYBR Green Master Mix on an ABI QuantStudio 5. Primers are described in Supplementary Table 3.

### Glucose transport

Glucose transport assays were performed as previously described^63^. Adipocytes were cultured in reduced media volumes as previously described for 48 h prior to the experiment (125 μl media per well of 48-well plate, media changed every 24 h) to improve insulin sensitivity^63^. Cells were serum-starved for 2 h in DMEM containing 0.2% BSA at 37°C, 10% CO_2_, washed and incubated in pre-warmed Krebs–Ringer phosphate (KRP) buffer containing 0.2% bovine serum albumin (KRP buffer; 0.6 mM Na_2_HPO_4_, 0.4 mM NaH_2_PO_4_, 120 mM NaCl, 6 mM KCl, 1 mM CaCl_2_, 1.2 mM MgSO_4_ and 12.5 mM HEPES (pH 7.4)) for 10 min, and then stimulated with 0.5 or 100 nM insulin for 20 min. To determine non-specific glucose uptake, 25 μM cytochalasin B (in ethanol, Sigma Aldrich) was added to control wells before addition of 2-[^3^H]deoxyglucose (2-DG) (PerkinElmer). During the final 5 min, 2-DG (0.25 μCi, 50 μM) was added to cells to measure steady-state rates of 2-DG uptake. Cells were then moved to ice, washed with ice-cold PBS, and solubilised in PBS containing 1% (v/v) Triton X-100. Tracer uptake was quantified by liquid scintillation counting on the TriCarb 2900TR (PerkinElmer) and data normalised for protein content.

### Immunofluorescence

*Cell surface immunofluorescence plate assays (HA-C3ORF18/GLUT4/TFR)* PM translocation of HA-C3ORF18, GLUT4 and TfR in 3T3-L1 adipocytes was assessed as previously described^54,63^, with minor modifications. Adipocytes were cultured in reduced media volumes as previously described for 48 h prior to the experiment (50 μl media per well of 96-well plate, media changed every 24 h) to increase insulin sensitivity^63^. Cells were serum starved for 2 h and stimulated with insulin at doses and for times indicated. Where inhibitors were used, cells were pre-treated with 200 nM Wortmannin (Sigma, W1628), 10 μM GDC-0068 (Selleck Chemicals S2808), MK2206 (MedChemExpress, HY-10358) or U0126 (Cell guidance systems, S106) for 15 min prior to insulin stimulation. Cells were washed by gently immersing the 96-well plates 10 times in beakers containing ice-cold PBS (five washes in each of two 1 L beakers containing PBS; all subsequent PBS washes were performed using this method). Plates were placed on ice and residual PBS was removed with a multichannel pipette. Cells were fixed with 4% paraformaldehyde (PFA) for 5 min on ice and 10 min at room temperature. PFA was quenched with 50 mM glycine in PBS for 10 min. Cells were blocked with 5% normal swine serum (NSS; Dako, X0901) in PBS for 30 min. Cells were then incubated with primary antibody solution (Supplementary Table 3) in 2% NSS in PBS for 1 h at room temperature, washed in PBS, and incubated with secondary antibody solution in 2% NSS in PBS for 1 h at room temperature in the dark. Cells were stored in PBS containing 2.5% DABCO, 10% glycerol and pH 8.5 as well as sealed and kept at 4 °C in the dark before imaging. Plates were equilibrated to room temperature for 30 min before imaging.

Plates were imaged on the Perkin Elmer Opera Phenix High Content Screening System and mid-section confocal images were obtained using a x20 water objective (NA 1.0), with 2-pixel binning. Excitation wavelengths and emission filters used were as follows: surface HA: 488 nm, 500–550 nm; endogenous surface TfR: 568 nm, 570-630; endogenous surface GLUT4: 647 nm, 650-760; and Hoechst: 405 nm, 435–480 nm. The microscope settings were automated and predefined before imaging the entire plate, meaning identical positions in each well were sampled across all wells and conditions without any human supervision. Nine positions towards the centre of each well were selected for imaging. Following the acquisition, positions across each plate and well were inspected at random to ensure proper seeding/staining for quality assurance. Mean fluorescence intensity was calculated to determine surface amount of protein.

Surface translocation of HA-C3ORF18 in L6 myoblasts and myotubes was performed as described above, with plates imaged using a x40 water objective with 40-50 fields of view per well used to quantify mean fluorescence intensity.

Surface translocation of HA-C3ORF18 in SGBS adipocytes was performed as described above with minor modifications. Cells serum starved for 4 h, stimulated, washed with PBS and incubated with primary antibody on ice for 2 h. Cells were washed in PBS, fixed with 4% PFA for 5 min on ice and 10 min at room temperature and secondary antibodies added as described above. Plates were imaged using a x40 water objective and 18 fields of view used to quantify mean fluorescence intensity in each well.

#### Colocalisation immunofluorescence (Opera Phenix)

Immunofluorescence to assess colocalisation of C3ORF18, GLUT4 and TfR was performed as described above with 0.1% saponin added to the blocking, primary and secondary antibody solutions to permeabilise cells. Plates were imaged on the Perkin Elmer Opera Phenix High Content Screening System and mid-section confocal images were obtained using a ×63 water objective. Colocalisation Threshold plugin in ImageJ Fiji v2.9.0 was used to compute Mander’s coefficient.

#### Colocalisation immunofluorescence (SP8)

Differentiated 3T3-L1 adipocytes were reseeded onto glass coverslips and fixed with 4% PFA, permeabilised and blocked with 0.1% saponin and 5 % NSS before incubation with primary antibodies (Supplementary Table 3) in 2% NSS, 0.1% saponin in PBS for 1 h at room temperature. Coverslips were washed with PBS and incubated with secondary antibodies and BODIPY^TM^ 493/503 (1:1000, Invitrogen, D3922) in 2% NSS in PBS for 1 h at room temperature. Imaging was performed using a Leica SP8 microscope with a x63 Plan-Apochromat oil objective. Image acquisition was performed using Leica Application Software (Leica, v3.5.7.2) and images noise reduced using the median module with 2 iterations.

### Statistical analysis

All statistical analyses were performed in Graphpad Prism 10 unless otherwise stated. Two-tailed paired Student’s tests were used to compare the means between 2 groups. One/two-way ANOVA with Šidák correction for multiple comparisons was used for multigroup comparisons. A mixed-effects model was used for multiple comparisons in Extended Data Fig. 5c. Variations among replicates were expected to have normal distributions and equal variance.

## Acknowledgements

We thank Professor Martin Wabitsch (Division of Pediatric Endocrinology and Diabetes, University Medical Center Ulm, Germany) for generously providing SGBS cells and Professor David James (University of Sydney) for kindly providing 3T3-L1 and L6 cells. We also thank Mike Deery (Cambridge Centre for Proteomics) for performing the mass spectrometry analysis and Yagnesh Umarania (Cambridge Centre for Proteomics) for assistance with proteomic data processing. This work was supported by the Cell Imaging Facility at the Institute of Metabolic Science, who were funded by the Medical Research Council [MC_UU_00039].

## Funding

This work was supported by a MRC Career Development award (MR/S007091/1) and Project grant (MR/Z504592/1) awarded to D.J.F. O.J.C. was supported by a Wellcome Trust PhD studentship. J.A.C. was supported by a BBSRC iCASE award with AstraZeneca (BB/R505304/1). L.M.B. was supported by EU Horizon 2020 program INFRAIA project EPIC-XS (project 823839). D.C.G. was supported by a Biotechnology and Biological Sciences Research Council project grant (BB/W005905/1) and a Wellcome Trust/Royal Society Sir Henry Dale Fellowship (210481). D.B.S. is supported by the Wellcome Trust (WT 219417), the MRC (MR/X00970X/1), and The National Institute for Health Research (NIHR) Cambridge Biomedical Research Centre and NIHR Rare Disease Translational Research Collaboration.

## Author contributions

O.J.C.; Conceptualisation; Data Curation; Formal Analysis; Funding Acquisition; Investigation; Methodology; Project Administration; Software; Visualization; Writing – Original Draft Preparation; Writing – Review & Editing. J.A.C.; Data Curation; Formal Analysis; Funding Acquisition; Investigation; Writing – Review & Editing. L.M.B.; Data Curation; Formal Analysis; Funding Acquisition; Software; Writing – Review & Editing. H.L.; Data Curation; Formal Analysis; Investigation; Writing – Review & Editing. D.M.; Investigation; Writing – Review & Editing. L.L.; Investigation; Writing – Review & Editing. S.P.; Methodology; Supervision; Writing – Review & Editing. D.C.G.; Funding Acquisition; Supervision; Writing – Review & Editing. D.B.S.; Funding Acquisition; Supervision; Writing – Review & Editing. M.P.W.; Funding Acquisition; Supervision; Writing – Review & Editing. K.S.L.; Funding Acquisition; Supervision; Writing – Review & Editing. D.J.F.; Conceptualisation; Funding Acquisition; Investigation; Methodology; Project Administration; Resources; Supervision; Validation; Writing – Original Draft Preparation; Writing – Review & Editing.

## Conflicts of interest

None to declare.

## Notes

### Competing Interest Statement

The authors have declared no competing interest.

## References

1. Humphrey, S. J. et al. Dynamic adipocyte phosphoproteome reveals that Akt directly regulates mTORC2. Cell Metab 17, 1009–1020 (2013).

2. Menon, S. et al. Spatial control of the TSC complex integrates insulin and nutrient regulation of mTORC1 at the lysosome. Cell 156, 771–785 (2014).

3. Fazakerley, D. J., Koumanov, F. & Holman, G. D. GLUT4 On the move. Biochem J 479, 445–462 (2022).

4. Davis, R. J., Corvera, S. & Czech, M. P. Insulin stimulates cellular iron uptake and causes the redistribution of intracellular transferrin receptors to the plasma membrane. J Biol Chem 261, 8708–8711 (1986).

5. Tanner, L. I. & Lienhard, G. E. Insulin elicits a redistribution of transferrin receptors in 3T3-L1 adipocytes through an increase in the rate constant for receptor externalization. J Biol Chem 262, 8975–8980 (1987).

6. Oka, Y., Mottola, C., Oppenheimer, C. L. & Czech, M. P. Insulin activates the appearance of insulin-like growth factor II receptors on the adipocyte cell surface. Proc Natl Acad Sci U S A 81, 4028–4032 (1984).

7. Kandror, K. V. & Pilch, P. F. The insulin-like growth factor II/mannose 6-phosphate receptor utilizes the same membrane compartments as GLUT4 for insulin-dependent trafficking to and from the rat adipocyte cell surface. J Biol Chem 271, 21703–21708 (1996).

8. Prior, M. J. et al. Quantitative proteomic analysis of the adipocyte plasma membrane. J Proteome Res 10, 4970–4982 (2011).

9. van Gerwen, J., Shun-Shion, A. S. & Fazakerley, D. J. Insulin signalling and GLUT4 trafficking in insulin resistance. Biochem Soc Trans 51, 1057–1069 (2023).

10. Fazakerley, D. J. et al. Phosphoproteomics reveals rewiring of the insulin signaling network and multi-nodal defects in insulin resistance. Nat Commun 14, 923 (2023).

11. Fazakerley, D. J. et al. Proteomic Analysis of GLUT4 Storage Vesicles Reveals Tumor Suppressor Candidate 5 (TUSC5) as a Novel Regulator of Insulin Action in Adipocytes. J Biol Chem 290, 23528–23542 (2015).

12. Geladaki, A. et al. Combining LOPIT with differential ultracentrifugation for high-resolution spatial proteomics. Nat Commun 10, 331 (2019).

13. Weekes, M. P. et al. Proteomic plasma membrane profiling reveals an essential role for gp96 in the cell surface expression of LDLR family members, including the LDL receptor and LRP6. J Proteome Res 11, 1475–1484 (2012).

14. Crook, O. M. et al. Inferring differential subcellular localisation in comparative spatial proteomics using BANDLE. Nat Commun 13, 5948 (2022).

15. Gatto, L., Breckels, L. M., Wieczorek, S., Burger, T. & Lilley, K. S. Mass-spectrometry-based spatial proteomics data analysis using pRoloc and pRolocdata. Bioinformatics 30, 1322–1324 (2014).

16. Klingelhuber, F. et al. A spatiotemporal proteomic map of human adipogenesis. Nat Metab 6, 861–879 (2024).

17. Akil, A. et al. Septin 9 induces lipid droplets growth by a phosphatidylinositol-5-phosphate and microtubule-dependent mechanism hijacked by HCV. Nature Commun 7, 12203 (2016).

18. Song, P. X. et al. Septin 9 and phosphoinositides regulate lysosome localization and their association with lipid droplets. iScience 25, 104288 (2022).

19. Moreno-Castellanos, N. et al. The cytoskeletal protein septin 11 is associated with human obesity and is involved in adipocyte lipid storage and metabolism. Diabetologia 60, 324–335 (2017).

20. Chen, F. et al. FIT2 organizes lipid droplet biogenesis with ER tubule-forming proteins and septins. J Cell Biol 220, (2021).

21. Xu, L. et al. Adipocyte Septin-7 attenuates obesogenic adipogenesis and promotes lipolysis to prevent obesity. Mol Metab 102114 (2025).

22. Zhong, J. et al. adiposetissue.org: A knowledge portal integrating clinical and experimental data from human adipose tissue. Cell Metab (2025) doi:10.1016/j.cmet.2025.01.012.

23. Petrus, P. et al. Transforming Growth Factor-β3 Regulates Adipocyte Number in Subcutaneous White Adipose Tissue. Cell Rep 25, 551–560.e5 (2018).

24. Kerr, A. G., Andersson, D. P., Rydén, M., Arner, P. & Dahlman, I. Long-term changes in adipose tissue gene expression following bariatric surgery. J Intern Med 288, 219–233 (2020).

25. Arner, P., Andersson, D. P., Bäckdahl, J., Dahlman, I. & Rydén, M. Weight Gain and Impaired Glucose Metabolism in Women Are Predicted by Inefficient Subcutaneous Fat Cell Lipolysis. Cell Metab 28, 45–54.e3 (2018).

26. Keller, M. et al. Genome-wide DNA promoter methylation and transcriptome analysis in human adipose tissue unravels novel candidate genes for obesity. Mol Metab 6, 86–100 (2017).

27. Arner, E. et al. Adipose tissue microRNAs as regulators of CCL2 production in human obesity. Diabetes 61, 1986–1993 (2012).

28. Stancáková, A. et al. Hyperglycemia and a common variant of GCKR are associated with the levels of eight amino acids in 9,369 Finnish men. Diabetes 61, 1895–1902 (2012).

29. Raulerson, C. K. et al. Adipose Tissue Gene Expression Associations Reveal Hundreds of Candidate Genes for Cardiometabolic Traits. Am J Hum Genet 105, 773–787 (2019).

30. Civelek, M. et al. Genetic Regulation of Adipose Gene Expression and Cardio-Metabolic Traits. Am J Hum Genet 100, 428–443 (2017).

31. Krieg, L. et al. Multiomics reveal unique signatures of human epiploic adipose tissue related to systemic insulin resistance. Gut 71, 2179–2193 (2022).

32. Arner, P. et al. The epigenetic signature of systemic insulin resistance in obese women. Diabetologia 59, 2393–2405 (2016).

33. Imbert, A. et al. Network Analyses Reveal Negative Link Between Changes in Adipose Tissue GDF15 and BMI During Dietary-induced Weight Loss. J Clin Endocrinol Metab 107, e130–e142 (2022).

34. Armenise, C. et al. Transcriptome profiling from adipose tissue during a low-calorie diet reveals predictors of weight and glycemic outcomes in obese, nondiabetic subjects. Am J Clin Nutr 106, 736–746 (2017).

35. Winnier, D. A. et al. Transcriptomic identification of ADH1B as a novel candidate gene for obesity and insulin resistance in human adipose tissue in Mexican Americans from the Veterans Administration Genetic Epidemiology Study (VAGES). PLoS One 10, e0119941 (2015).

36. Nono Nankam, P. A., et al. Distinct abdominal and gluteal adipose tissue transcriptome signatures are altered by exercise training in African women with obesity. Sci Rep 10, 10240 (2020).

37. Vink, R. G. et al. Adipose tissue gene expression is differentially regulated with different rates of weight loss in overweight and obese humans. Int J Obes (Lond) 41, 309–316 (2017).

38. Salcedo-Tacuma, D. et al. Transcriptome dataset of omental and subcutaneous adipose tissues from gestational diabetes patients. Sci Data 9, 344 (2022).

39. MacLaren, R. E., Cui, W., Lu, H., Simard, S. & Cianflone, K. Association of adipocyte genes with ASP expression: a microarray analysis of subcutaneous and omental adipose tissue in morbidly obese subjects. BMC Med Genomics 3, 3 (2010).

40. Johansson, L. E. et al. Differential gene expression in adipose tissue from obese human subjects during weight loss and weight maintenance. Am J Clin Nutr 96, 196–207 (2012).

41. Matualatupauw, J. C., Bohl, M., Gregersen, S., Hermansen, K. & Afman, L. A. Dietary medium-chain saturated fatty acids induce gene expression of energy metabolism-related pathways in adipose tissue of abdominally obese subjects. Int J Obes (Lond) 41, 1348–1354 (2017).

42. du Plessis, J. et al. Association of Adipose Tissue Inflammation With Histologic Severity of Nonalcoholic Fatty Liver Disease. Gastroenterology 149, 635–48.e14 (2015).

43. Van Bussel, I. P. G. et al. The impact of protein quantity during energy restriction on genome-wide gene expression in adipose tissue of obese humans. Int J Obes (Lond) 41, 1114–1120 (2017).

44. James, D. E., Brown, R., Navarro, J. & Pilch, P. F. Insulin-regulatable tissues express a unique insulin-sensitive glucose transport protein. Nature 333, 183–185 (1988).

45. Ross, S. A. et al. Characterization of the insulin-regulated membrane aminopeptidase in 3T3-L1 adipocytes. J Biol Chem 271, 3328–3332 (1996).

46. Keller, S. R., Scott, H. M., Mastick, C. C., Aebersold, R. & Lienhard, G. E. Cloning and characterization of a novel insulin-regulated membrane aminopeptidase from Glut4 vesicles. J Biol Chem 270, 23612–23618 (1995).

47. Larance, M. et al. Characterization of the role of the Rab GTPase-activating protein AS160 in insulin-regulated GLUT4 trafficking. J Biol Chem 280, 37803–37813 (2005).

48. Jedrychowski, M. P. et al. Proteomic analysis of GLUT4 storage vesicles reveals LRP1 to be an important vesicle component and target of insulin signaling. J Biol Chem 285, 104–114 (2010).

49. Yang, J. & Holman, G. D. Comparison of GLUT4 and GLUT1 subcellular trafficking in basal and insulin-stimulated 3T3-L1 cells. J Biol Chem 268, 4600–4603 (1993).

50. Cushman, S. W. & Wardzala, L. J. Potential mechanism of insulin action on glucose transport in the isolated rat adipose cell. Apparent translocation of intracellular transport systems to the plasma membrane. J Biol Chem 255, 4758–4762 (1980).

51. Suzuki, K. & Kono, T. Evidence that insulin causes translocation of glucose transport activity to the plasma membrane from an intracellular storage site. Proc Natl Acad Sci U S A 77, 2542–2545 (1980).

52. Wardzala, L. J., Cushman, S. W. & Salans, L. B. Mechanism of insulin action on glucose transport in the isolated rat adipose cell. Enhancement of the number of functional transport systems. J Biol Chem 253, 8002–8005 (1978).

53. Simpson, I. A., Hedo, J. A. & Cushman, S. W. Insulin-induced internalization of the insulin receptor in the isolated rat adipose cell. Detection of both major receptor subunits following their biosynthetic labeling in culture. Diabetes 33, 13–18 (1984).

54. Diaz-Vegas, A. et al. A high-content endogenous GLUT4 trafficking assay reveals new aspects of adipocyte biology. Life Sci Alliance 6, (2023).

55. Clarke, J. F., Young, P. W., Yonezawa, K., Kasuga, M. & Holman, G. D. Inhibition of the translocation of GLUT1 and GLUT4 in 3T3-L1 cells by the phosphatidylinositol 3-kinase inhibitor, wortmannin. Biochem J 300 ( **Pt 3**), 631–635 (1994).

56. Justice, A. E. et al. Protein-coding variants implicate novel genes related to lipid homeostasis contributing to body-fat distribution. Nat Genet 51, 452–469 (2019).

57. Gilleron, J., Gerdes, J. M. & Zeigerer, A. Metabolic regulation through the endosomal system. Traffic 20, 552–570 (2019).

58. Dai, X. et al. AMPK-dependent phosphorylation of the GATOR2 component WDR24 suppresses glucose-mediated mTORC1 activation. Nat Metab 5, 265–276 (2023).

59. Tews, D. et al. 20 Years with SGBS cells - a versatile in vitro model of human adipocyte biology. Int J Obes (Lond) 46, 1939–1947 (2022).

60. Wabitsch, M. et al. Characterization of a human preadipocyte cell strain with high capacity for adipose differentiation. Int J Obes Relat Metab Disord 25, 8–15 (2001).

61. Govers, R., Coster, A. C. F. & James, D. E. Insulin increases cell surface GLUT4 levels by dose dependently discharging GLUT4 into a cell surface recycling pathway. Mol Cell Biol 24, 6456–6466 (2004).

62. Morita, S., Kojima, T. & Kitamura, T. Plat-E: an efficient and stable system for transient packaging of retroviruses. Gene Ther 7, 1063–1066 (2000).

63. Tan, J. et al. Limited oxygen in standard cell culture alters metabolism and function of differentiated cells. EMBO J 43, 2127–2165 (2024).

64. Brasaemle, D. L. & Wolins, N. E. Isolation of Lipid Droplets from Cells by Density Gradient Centrifugation. Curr Protoc Cell Biol 72, 3.15.1–3.15.13 (2016).

65. Yu, G., Wang, L.-G., Han, Y. & He, Q.-Y. clusterProfiler: an R package for comparing biological themes among gene clusters. OMICS 16, 284–287 (2012).

66. Wu, T. et al. clusterProfiler 4.0: A universal enrichment tool for interpreting omics data. Innovation (Camb) 2, 100141 (2021).

67. Gatto, L. & Lilley, K. S. MSnbase-an R/Bioconductor package for isobaric tagged mass spectrometry data visualization, processing and quantitation. Bioinformatics 28, 288– 289 (2012).

68. Breckels, L. M., Mulvey, C. M., Lilley, K. S. & Gatto, L. A Bioconductor workflow for processing and analysing spatial proteomics data. F1000Res 5, 2926 (2016).

69. Mulvey, C. M. et al. Using hyperLOPIT to perform high-resolution mapping of the spatial proteome. Nat Protoc 12, 1110–1135 (2017).

70. Itzhak, D. N., Tyanova, S., Cox, J. & Borner, G. H. Global, quantitative and dynamic mapping of protein subcellular localization. Elife 5, (2016).

71. Crook O, B. L. bandle: An R package for the Bayesian analysis of differential subcellular localisation experiments. R package version 1.10.0. http://github.com/ococrook/bandle (2024).

72. *bandle: An R package for the Bayesian analysis of differential subcellular localisation experiments*. https://bioconductor.org/packages/release/bioc/html/bandle.html (2024).

73. Gene Ontology Consortium. The Gene Ontology resource: enriching a GOld mine. Nucleic Acids Res 49, D325–D334 (2021).

74. Kanehisa, M., Furumichi, M., Sato, Y., Kawashima, M. & Ishiguro-Watanabe, M. KEGG for taxonomy-based analysis of pathways and genomes. Nucleic Acids Res 51, D587– D592 (2023).

75. Milacic, M. et al. The Reactome Pathway Knowledgebase 2024. Nucleic Acids Res 52, D672–D678 (2024).

76. Benjamini, Y. & Hochberg, Y. Controlling the false discovery rate: a practical and powerful approach to multiple testing. Journal of the Royal Statistical Society: series B (Methodological) 57, 289–300 (1995).

77. McAlister, G. C. et al. Increasing the multiplexing capacity of TMTs using reporter ion isotopologues with isobaric masses. Anal Chem 84, 7469–7478 (2012).

78. McAlister, G. C. et al. MultiNotch MS3 enables accurate, sensitive, and multiplexed detection of differential expression across cancer cell line proteomes. Anal Chem 86, 7150–7158 (2014).

79. Huttlin, E. L. et al. A tissue-specific atlas of mouse protein phosphorylation and expression. Cell 143, 1174–1189 (2010).

80. Cox, J. & Mann, M. MaxQuant enables high peptide identification rates, individualized p.p.b.-range mass accuracies and proteome-wide protein quantification. Nature Biotechnology 26, 1367–1372 (2008).

81. Williamson, A. et al. Genome-wide association study and functional characterization identifies candidate genes for insulin-stimulated glucose uptake. Nat Genet 55, 973–983 (2023).

82. Pereira, C. et al. The exocyst complex is an essential component of the mammalian constitutive secretory pathway. J Cell Biol 222, (2023).

