## Extended Data Fig. 1-5 for "Dynamic subcellular proteomics identifies novel regulators of adipocyte insulin action"

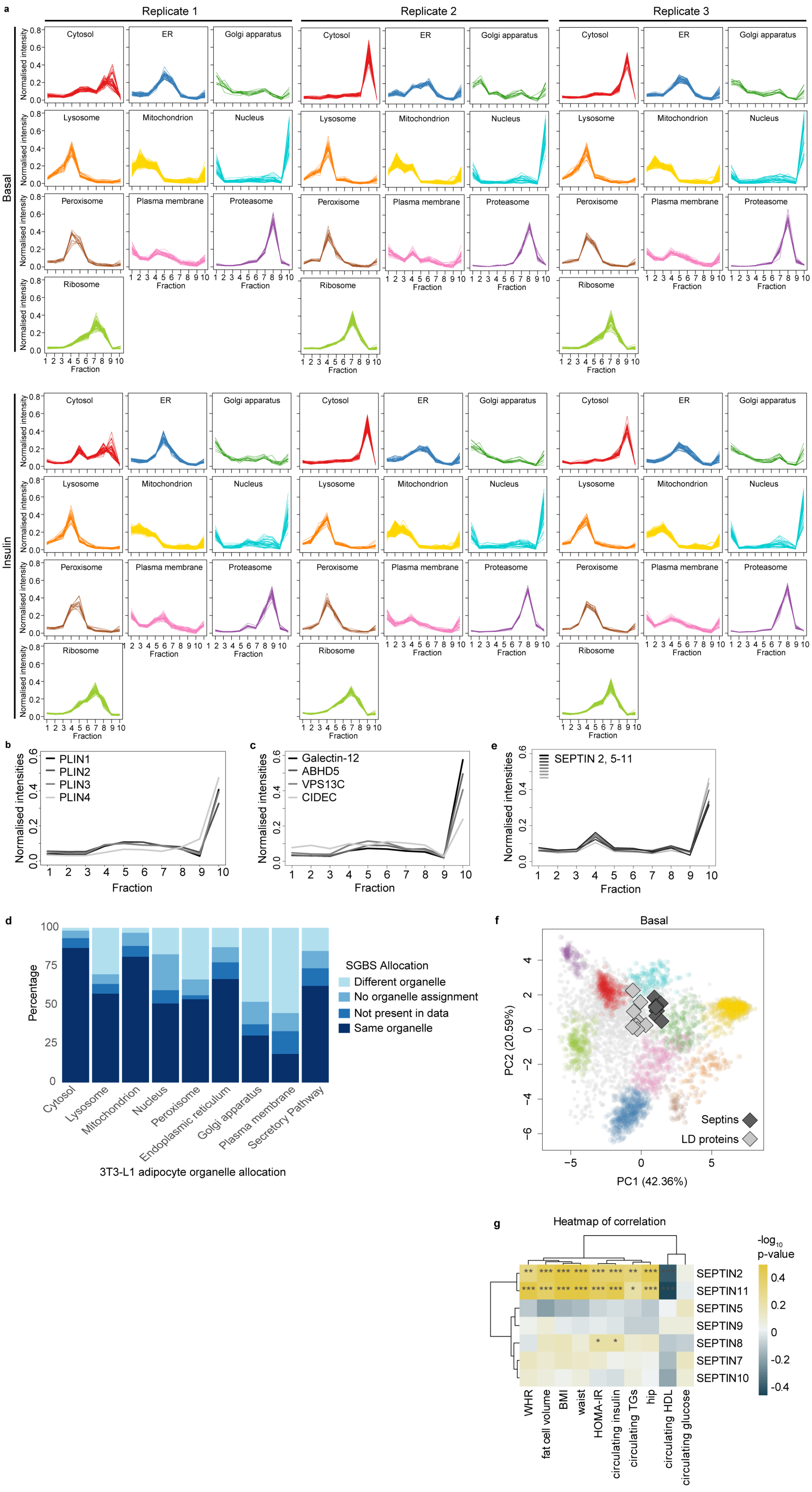
a
Replicate 1
Replicate 2
Replicate 3
Basal
Insulin
Normalised intensity
Cytosol
ER
Golgi apparatus
Lysosome
Mitochondrion
Nucleus
Peroxisome
Plasma membrane
Proteasome
Ribosome
Fraction
b
PLIN1
PLIN2
PLIN3
PLIN4
Normalised intensities
Fraction
c
Galectin-12
ABHD5
VPS13C
CIDEC
Normalised intensities
Fraction
e
SEPTIN 2, 5-11
Normalised intensities
Fraction
d
Percentage
SGBS Allocation
Different organelle
No organelle assignment
Not present in data
Same organelle
Cytosol
Lysosome
Mitochondrion
Nucleus
Peroxisome
Endoplasmic reticulum
Golgi apparatus
Plasma membrane
Secretory Pathway
3T3-L1 adipocyte organelle allocation
f
Basal
Septins
LD proteins
PC1 (42.36%)
PC2 (20.59%)
g
Heatmap of correlation
-log10 p-value
SEPTIN2
SEPTIN11
SEPTIN5
SEPTIN9
SEPTIN8
SEPTIN7
SEPTIN10
WHR
fat cell volume
BMI
waist
HOMA-IR
circulating insulin
circulating TGs
hip
circulating HDL
circulating glucose

**Extended Figure 1**. a) Distribution of marker proteins (n = 531) across fractions in each replicate. Mean distribution of (b) perilipin proteins, (c) other known lipid droplet associated proteins and (e) septin proteins identified (septin 2, 5-12) across fractions under basal conditions (n = 3). d) Comparison of organelle allocation of proteins present in 3T3-L1 adipocytes to allocation in SGBS adipocytes[16]. f) PCA projection of basal LOPIT-DC map with septins (2, 5-11, dark grey diamonds) and lipid droplet marker proteins (PLIN1-4, CIDEC, VPS13C, galectin-12, ABHD5, light grey diamonds) highlighted. g) Spearman's correlation of septin protein expression in omental adipose with metabolic clinical features. Figure generated from adiposetissue.org[22] using data from [23–43] (*** pFDR <0.001, ** pFDR < 0.01, * pFDR < 0.05).

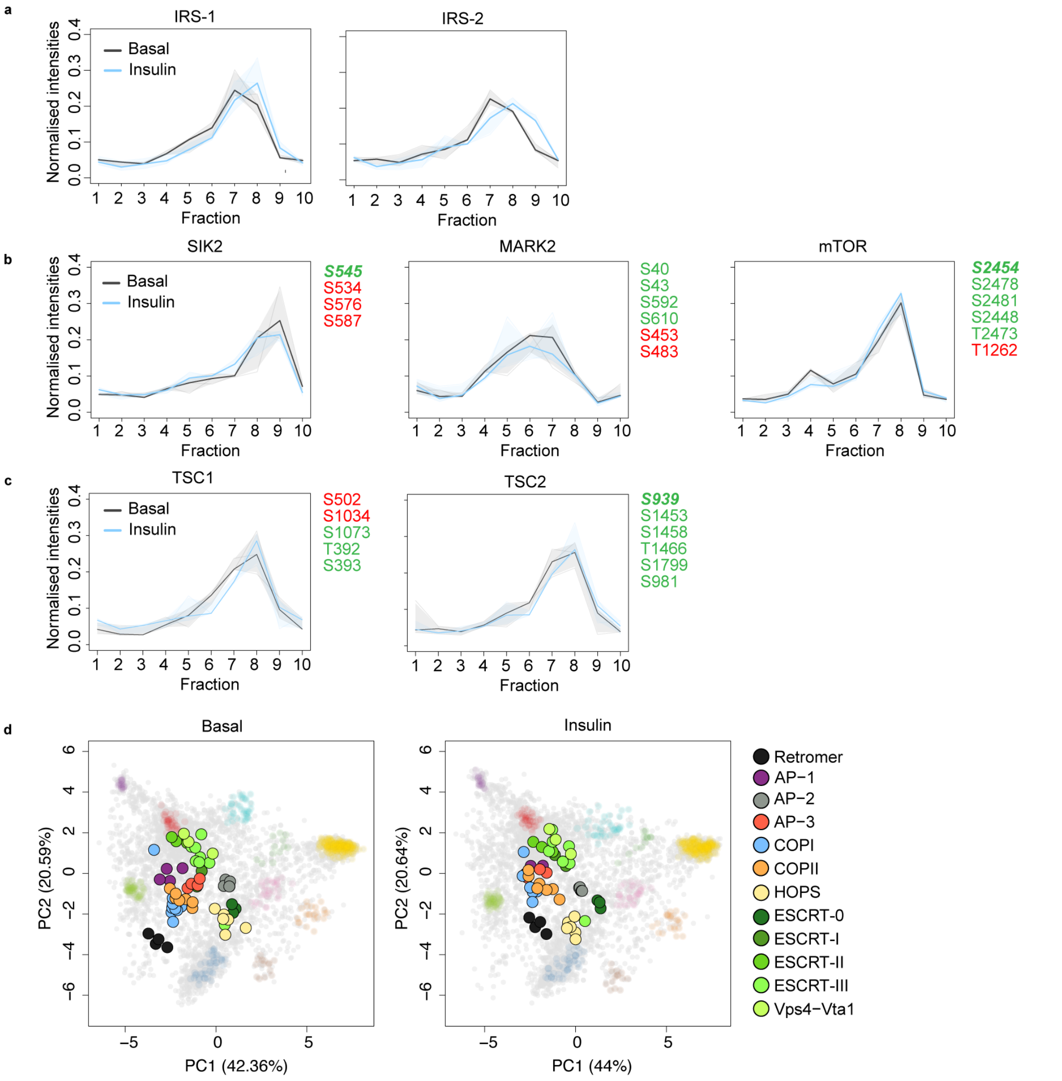


**Extended Figure 2:** Distribution of (a) IRS-1 and IRS-2, (b) SIK2, MARK2 and mTOR, and (c) TSC1/2 subunits across fractions in basal (grey) and insulin-treated (blue) adipocytes (solid line = mean, shaded area 95% CI, n = 3 per condition), and insulin-regulated phosphorylation sites reported in Fazakerley (2023)[10] and Humphrey (2013)[1] (green = increased and red = decreased phosphorylation, regular font = found in either Fazakerley and Humphrey, bold italic = found in both Fazakerley and Humphrey). IRS-1/2 have >20 regulated phosphosites and are listed in Supplementary Table 1. d) PCA projection of concatenated basal (left) and insulin (right) LOPIT-DC data with cytosolic cargo sorting complexes highlighted. Only subunits found in all 6 replicates are shown. Retromer - VPS26A, VPS26B, VPS29, VPS35; ESCRT-0 - HRS, STAM2; ESCRT-I - VPS23, VPS37C; ESCRT-II - SNF8, VPS25, VPS36; ESCRT-III - CHMP1A, CHMP2A, CHMP2B, CHMP3, CHMP4B, CHMP5, CHMP6, IST1; Vps4-Vta1 - VPS4A, VPS4B, VTA1; AP-1 - AP1B1, AP1G1, AP1M1, AP1S1; AP-2 - AP2A1, AP2A2, AP2S1, AP2M1, AP2B1; AP-3 - AP3B1, AP3D1, AP3M1, AP3S1; COPI - COPA, COPB1, COPB2, COPE, COPG1, COPG2, COPZ1, COPZ2, ARCN1; COPII - SAR1A, SAR1B, SEC13, SEC23A, SEC23B, SEC24A, SEC31A; HOPS - VPS11, VPS16, VPS18, VPS33A, VPS39, VPS41.

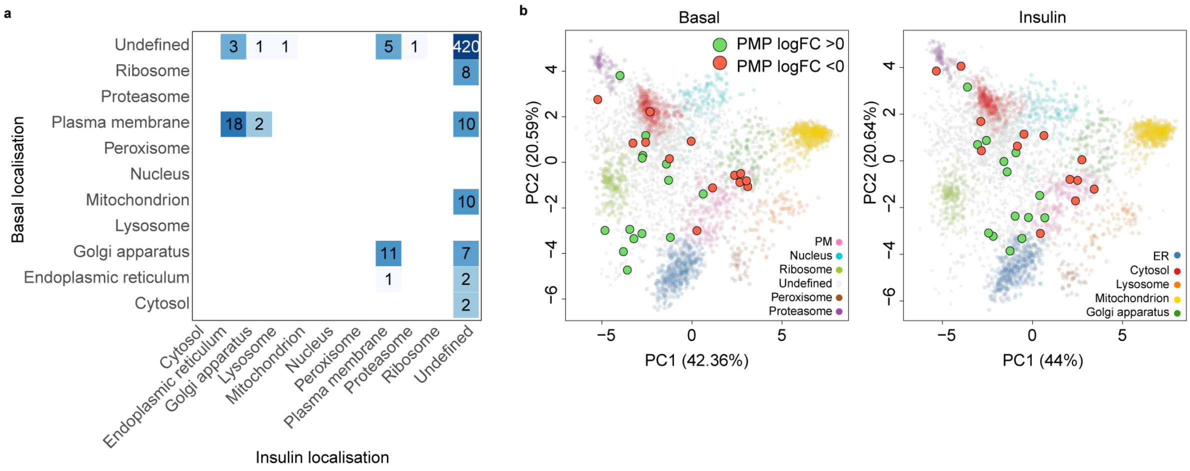


**Extended Figure 3:** a) Organelle assignment of proteins predicted to relocalise with high-confidence (*diff. loc. prob*. = 1) under basal and insulin-treated conditions. b) PCA projection of LOPIT-DC adipocyte subcellular map under basal (left) and insulin-stimulated (right) conditions with proteins predicted to move with high-confidence (*diff. loc. prob*. = 1) also up (green) or downregulated (red) ($p$<0.05) in plasma membrane proteomics (PMP) data highlighted.

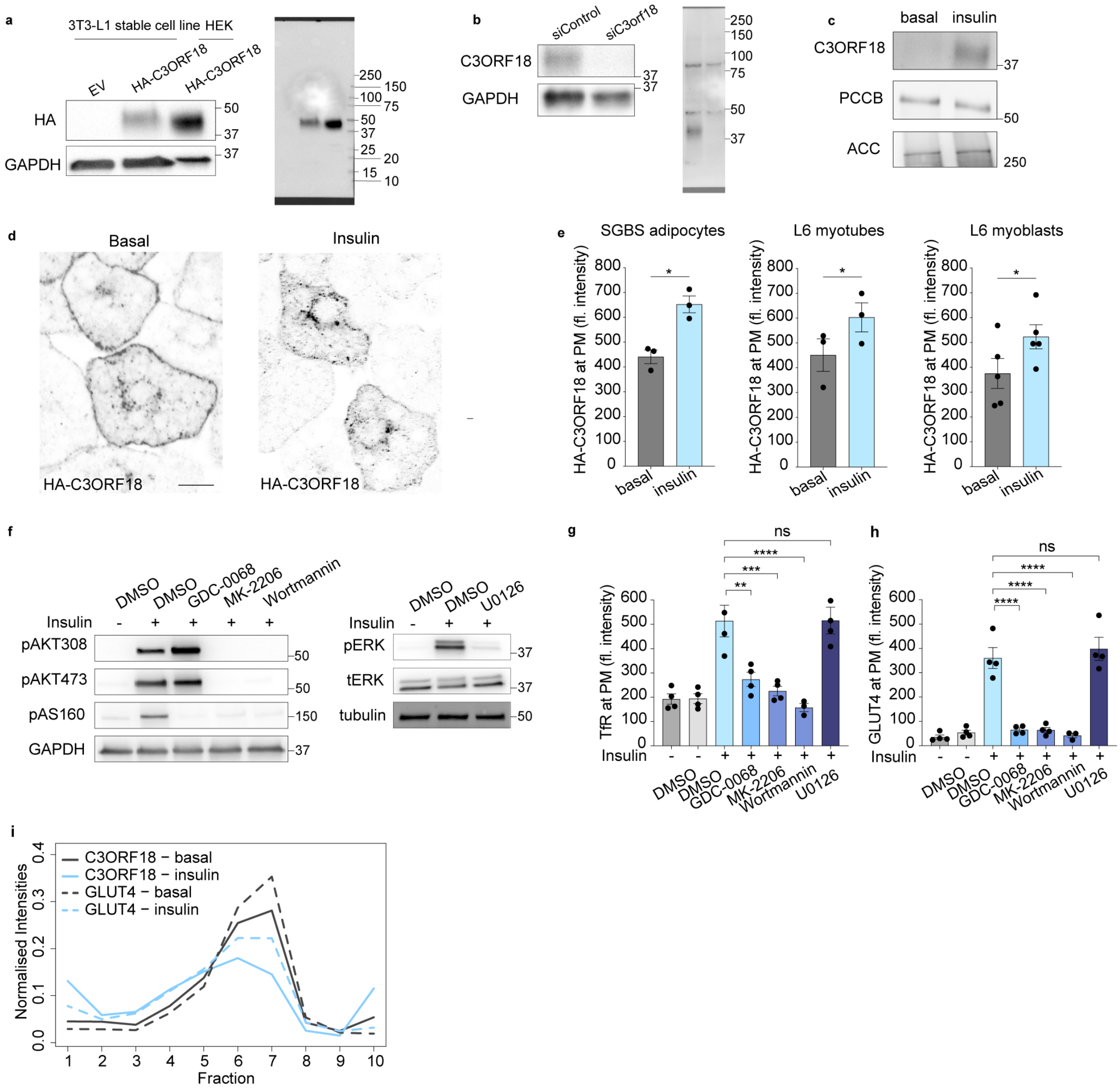


**Extended Figure 4:** a) Western blot analysis of HA expression in 3T3-L1 adipocytes stable cell lines expressing HA-C3ORF18 or the empty control vector (EV), and HEK-293 cells transiently overexpressing HA-C3ORF18. 4-20% SDS-PAGE gel used to resolve proteins, full membrane shown on right. b) Western blot analysis of C3ORF18 expression in 3T3-L1 adipocytes with siRNA-mediated depletion of C3ORF18. 10% SDS-PAGE gel used to resolve proteins, full membrane shown on right. c) Western blot analysis of C3ORF18 in the PM fraction of 3T3-L1 adipocytes isolated for plasma membrane proteomics as described in Fig. 3b. d) Immunostaining of HA-C3ORF18 overexpressing 3T3-L1 adipocytes under basal and insulin-stimulated (100 nM, 30 min) conditions (scale bar = 10 µm). e) Relative fluorescence intensity of anti-HA surface staining in basal and insulin-stimulated HA-C3ORF18 expressing SGBS adipocytes, L6 myoblasts and myotubes (n = 3-6 biological replicates). f) Western blot analysis of 3T3-L1 adipocytes treated with AKT-PI3K (GDC-0068, MK-2206, Wortmannin) or MAPK signalling inhibitors (U0126) for 15 min prior to insulin stimulation (100 nM, 30 min). Representative blot shown, n = 3. Relative fluorescence intensity of surface TfR (g) and GLUT4 (h) in 3T3-L1 adipocytes treated with AKT-PI3K (GDC-0068, MK-2206, Wortmannin) or MAPK signalling inhibitors (U0126) prior to insulin stimulation (100 nM, 30 min, n = 3-4 biological replicates). i) Mean distribution of C3ORF18 (solid lines) and GLUT4 (dashed lines) across subcellular fractions under basal (grey) and insulin-stimulated (blue) conditions. All data represented as mean ± SEM (e, g, h). n.s. non-significant; *$p < 0.05$; **$p < 0.01$; ***$p < 0.01$; ****$p < 0.0001$ by paired two-tailed Student's *t*-test (e) or one-way ANOVA with Šidák's multiple comparisons test (g, h).

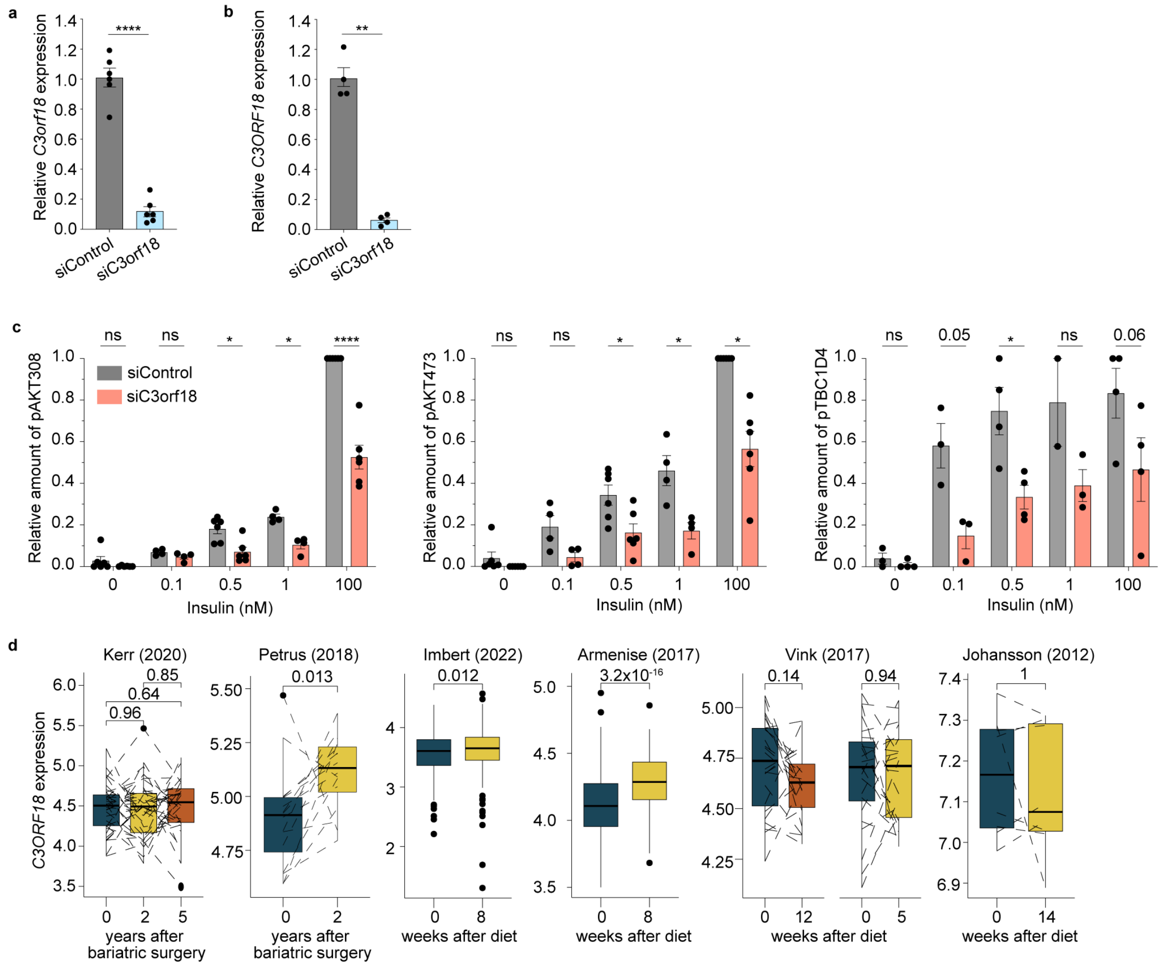


**Extended Figure 5:** Relative expression of *C3ORF18* in (a) 3T3-L1 and (b) SGBS adipocytes after 8 d knockdown post-differentiation (n = 4-6 biological replicates). c) Quantification of pT308AKT, pS473AKT and pT642TBC1D4 in 3T3-L1 adipocytes following C3ORF18 depletion, stimulated with 0, 0.1, 0.5, 1 or 100 nM insulin for 30 min treated with 0, 0.1, 0.5, 1 or 100 nM insulin for 30 min. Normalised to GAPDH (n = 4-6 biological replicates, representative blot shown in Fig. 5e). d) *C3ORF18* mRNA expression in adipose tissue following weight loss. Figure generated from adiposetissue.org[22] using data from [23–43] and analysed using Wilcoxon's matched pairs test. All other data represented as mean ± SEM (a-c). n.s. non-significant; *$p < 0.05$; **$p < 0.01$; ****$p < 0.0001$ by paired two-tailed Student's *t*-test (a,b) or by a mixed effects model with Šidák's multiple comparisons test (c).
